# Selection for high metabolic rate reduces gut microbiota responsiveness to dietary restriction in bank voles

**DOI:** 10.64898/2026.09.18.752420

**Authors:** Joanna K. Baran, Paweł Koteja, Tea Ivancic, Kunika, Małgorzata M. Lipowska, Piotr Łukasik, Edyta T. Sadowska, Phillip C. Watts, Anni M. Hämäläinen

## Abstract

Host-microbiota interactions are crucial in adapting to environmental changes, such as fluctuating energy availability. Yet, the extent to which hosts and their microbiota exhibit coordinated plasticity in response to transient energy limitation remains unclear. Here, we examined the capacity for parallel reversible plasticity in hosts and gut microbiota to temporary dietary restriction in hosts with differing energy needs. We subjected bank voles (*Clethrionomys glareolus*) from a long-term artificial selection experiment for high aerobic metabolism (A-lines) and control lines (C-lines) to dietary restriction (food dilution with insoluble fiber) followed by a recovery period. We measured traits relevant for vole energy consumption and fecal microbiota composition at baseline, after dietary restriction, and after recovery. We hypothesized that A-line voles would experience more severe and prolonged physiological effects of dietary restriction than C-line voles, which might be counteracted by higher microbiota responsiveness.

Dietary restriction caused a temporary increase in food consumption, decrease in body mass and resting metabolic rate, and significant shifts in gut microbiota composition, all of which returned to near baseline levels during the recovery period. These effects were largely similar for the selection lines, but contrary to our prediction, the effect of dietary restriction on bacterial diversity was stronger in C-lines. This suggests higher resistance of the A-line microbiota to dietary changes, perhaps resulting from increased host control over microbial communities due to the directional selection for high metabolic capacity. Overall, our findings highlight a substantial capacity for reversible plasticity in host physiology and gut microbiota in fluctuating environments.

## Introduction

Fluctuations in food availability are a typical challenge for animals in nature, requiring adaptive coping mechanisms to overcome energy and nutrient shortages. With its critical role in digesting complex foodstuffs, the gut microbiota may be particularly important in responding to changes in energy availability and expenditure (Kolodny and Schulenburg 2020; Voolstra and Ziegler 2020; Lindsay et al. 2020; Martin et al. 2007). The gut microbiota can also buffer the host against changes in energy availability by participating in the regulation of host energy metabolism. As the host and microbiota coevolve under shared environmental conditions, they may coadapt to facilitate optimal metabolism, energy usage, and degree of plasticity for their environment. In natural populations these interlinked processes are difficult to study, although adaptive modulation of energy harvest through microbiota changes is suggested by, for example, seasonal patterns in the microbiota of wild primates (Amato et al. 2015; Baniel et al. 2021). Yet, the dynamics of host metabolism and microbiota have rarely been examined experimentally in the context of energy availability fluctuation. In this study, we isolated the effects of the host energy needs and energy availability on the microbiota by applying temporary dietary restriction to bank voles *Clethrionomys* (*Myodes) glareolus* that had been artificially selected for differing metabolic rates.

Differences among animal individuals in metabolic rate contribute to their coping under varying energy availability conditions (Norin and Metcalfe 2019). Individuals with a fast basal metabolism and the associated higher maintenance energy needs may be more affected by reduced energy availability. Also, individuals vary in their capacity to adjust energy metabolism (level of metabolic plasticity) to environmental variation, such as changes in food availability (Glazier and Gjoni 2024). For example, dietary restriction induced a more significant reduction in metabolic rate or alteration in body composition in mice with a higher rate of basal metabolism (Brzęk et al. 2012). Differential sensitivity to energy shortage can be balanced by metabolic flexibility, the capacity to switch between different energy substrates, thus regulating energy consumption, storage, and mobilization in response to energy availability fluctuations (Smith et al. 2018).

In addition to host traits, gut microbiota has an important role in regulating the host energy balance. Gut microbiota have a well-documented role in energy harvest, including a significant role of microbiota composition in mediating host adiposity (Tremaroli and Bäckhed 2012; Heiss and Olofsson 2017). Strong evidence of this comes from fecal microbiota transplant studies showing that the metabolic phenotype can be transmitted by gut microbiota. For example, mice receiving microbiota from obese donors tend to develop obesity (Turnbaugh et al. 2006), but see also (Dalby 2023). When energy availability is low, gut microbiota may, in turn, facilitate host coping, for example through enrichment of bacteria that extract energy from the diet with higher efficiency (Amato et al. 2015; Baniel et al. 2021). The importance of the microbiota in adapting to dietary change is reinforced by the speed of the response: gut microbiota composition may change substantially within days of a dietary change (David et al. 2014; Turnbaugh et al. 2009). Moreover, such community changes are largely repeatable and reversible (E. M. Anderson et al. 2022; Low et al. 2021), although some shifts may persist over long time periods (Bourdeau-Julien et al. 2023).

The host metabolism and gut microbiota thus both play an important role in adapting to fluctuations in energy availability, with potentially significant fitness consequences. How the host metabolic phenotype interacts with the microbiota in response to dietary change remains poorly understood, however. Diet typically has a stronger effect on microbial community structure than host genotype (Carmody et al. 2015; Korach-Rechtman et al. 2019), although variation in numerous host genes has been linked to differences in microbiota composition (Spor et al. 2011; Wilde et al. 2024; Chaston et al. 2016; Bolnick et al. 2014; Salzman et al. 2010). The response of the microbiota to dietary intervention is shaped by the initial microbiota composition together with host metabolic characteristics and gene expression (Klimenko et al. 2022), but little is still known about host effects on microbiota flexibility, i.e., the capacity of microbial communities to respond to environmental change.

To examine the significance of the host metabolic background and the microbiota in responding to fluctuation in energy availability, we exposed bank voles with differing metabolic profiles to temporary energy limitation by diluting their food with cellulose and quantified longitudinal changes in host physiology and gut microbiota. We used animals from a long-term selection experiment, in which bank voles exposed to artificial selection for fast metabolism (A lines) have evolved a higher exercise-induced and basal metabolic rate and increased their daily food consumption compared with unselected control (C lines) voles (Sadowska et al. 2015b; Grosiak et al. 2020; Hseiky et al. 2026). A-line bank voles thus have higher energy needs to sustain their metabolism, and are expected to experience more adverse effects under conditions of limited energy availability. This difference in metabolic demand provides a useful framework for testing how hosts with contrasting energy requirements respond to energy limitation. Experimental calorie restriction influences the energy expenditure (Ramsey et al. 2000) and resting metabolic rate in, for example, rodents (Sohal et al. 2009; Faulks et al. 2006; Selman et al. 2005) and humans (Martin et al. 2007), and causes morphological changes in the gut (Naya et al. 2007). The role of microbiota in mediating these effects is increasingly recognized: dietary restriction influences microbiota composition, which can in turn affect host physiology and behavior (Wang et al. 2024). Here, we tested the general hypothesis that a greater metabolic need, caused either by host phenotype and/or an energy deficient diet, should elicit largely reversible changes in gut microbiota composition to allow the host to meet its energy needs. The study design and general predictions are shown in Fig. 1.

**Figure 1:**
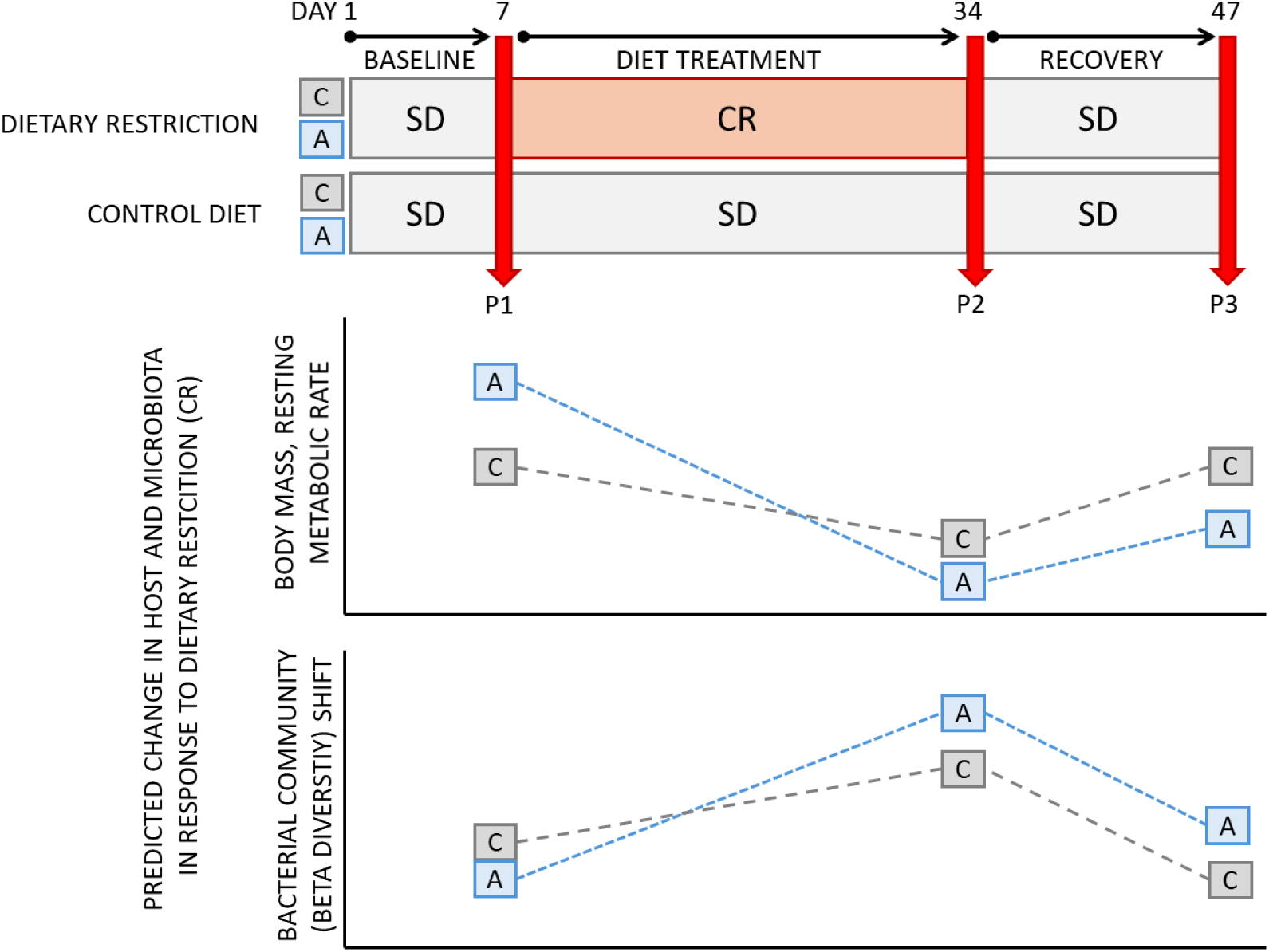
Study design, timeline, and *a priori* predictions. After a 6-day baseline period on a standard diet (SD; phase P1), bank voles from high-metabolism selection lines (A) and control lines (C) were assigned to either a standard diet (SD) or a cellulose-diluted dietary restriction treatment (CR) for 27 days (phase P2), followed by a 13-day recovery period on the standard diet (phase P3). Physiological traits and fecal samples for microbiota analyses were collected at the end of each phase (P1-P3). Animals were euthanized and dissected on day 48. Graphs illustrate the predicted effects of dietary restriction and recovery on host body mass, resting metabolic rate, and gut microbiota composition (β-diversity first distances) relative to baseline.

We posed two research questions and tested their associated hypotheses:

### 1. Do differences in animals’ energy needs influence their physiological and microbiota response to dietary restriction?

H1.1 We hypothesize that dietary restriction will elicit stronger physiological responses in A-line voles than in C-line voles, reflecting compensation for a larger energetic shortfall associated with their higher baseline energy requirements. Specifically, A-line voles are expected to show greater reductions in body mass and resting metabolic rate (RMR) under dietary restriction.

H1.2 We predict that food intake will differ between selection lines both before and during dietary restriction. At baseline, A-line voles are expected to consume more food than C-line voles, and under dietary restriction they should exhibit a greater relative increase in food intake as a compensatory response.

H1.3 If the gut microbiota buffers or mediates host physiological responses to energetic constraints, then dietary restriction should induce larger shifts in microbiota composition and diversity in A-line voles than in C-line voles, mirroring their stronger host-level responses.

### 2. Do differences in animals’ energy needs impact their capacity for reversible plasticity in response to dietary changes?

H2.1 We hypothesize that most physiological traits affected by dietary restriction—including body mass, food consumption, and RMR—will be reversible once the original diet is resumed. However, if dietary restriction imposes a stronger challenge on A-line voles, recovery of these traits may be slower or incomplete due to carry-over effects. If physiological traits do not return to baseline levels by the end of the recovery period, or if body composition and digestive tract size remain altered, this would indicate permanent or slowly reversible effects of dietary restriction rather than fully reversible plasticity.

H2.2 We predict partial to complete recovery of the gut microbiota following the recovery period. Any lingering effects of dietary restriction on microbial diversity or community composition are expected primarily in A-line voles if dietary restriction caused a larger initial microbiota shift in them.

Overall, our results indicate that exposing bank voles with differing energy needs to an energetic challenge caused pervasive yet remarkably reversible effects on both host physiology and the gut microbiota, while selection history exerted only subtle effects that were partly opposite to our predictions.

## Materials and Methods

The study design, animal husbandry, sampling protocols, measurement techniques and analytical approaches are described briefly below. Please refer to Supplementary Information (SI) for full methodological details.

### Animal model and selection experiment

This experiment was performed on laboratory-bred bank voles*, C. glareolus*, from generation 29 of a multidirectional artificial selection experiment described previously (Sadowska et al. 2015b, 2008). Briefly, the colony was founded in 2000 using ca. 320 wild-caught bank voles, and selection procedures were initiated after 5–6 generations of random breeding. Here, animals from high metabolism selection treatment lines (A), and unselected control lines (C) were used, with four replicate lines maintained within each line type. In the A-lines, selection was based on the maximum oxygen consumption rate per minute achieved during an 18-minute swimming trial at 38°C (V_O2, swim_), while C-lines were maintained as unselected control. By generation 27, voles from the A lines had an 84% higher V_O2, swim_ than those from the C lines (Jaromin et al. 2019), see also (Hanhimäki et al. 2022) and Fig. S1 in the SI.

A-line bank voles are also characterized by a higher basal metabolic rate (Sadowska et al. 2015b; Grosiak et al. 2020), thermogenic capacity (Dheyongera et al. 2016), aerobic capacity (forced-running V_O2max_) (Jaromin et al 2019), larger body mass (Grosiak et al. 2020; Lipowska et al. 2019), lower maximum corticosterone level (Lipowska et al. 2020), higher activity (Maiti et al. 2019), and higher daily food consumption rate (Dheyongera et al. 2016) compared to C-line animals. Additionally, the selection line types differ in the expression of several genes related to metabolic processes (Konczal 2015). In previous studies, no significant difference was found between the alpha or beta diversity of the gut microbiota (based on 16S amplicon sequencing) of A- and C-line bank voles under regular laboratory conditions (Kohl et al. 2016; Hanhimäki et al. 2022), but the response of the gut microbiota to environmental change differed somewhat between the line types (Hanhimäki et al. 2022).

Animals were maintained under standard conditions in dedicated animal facilities at the Jagiellonian University until enrollment into the experiment. Litters remained with their mothers until weaning (age 17 days), thus exposed to the maternal microbiota through birth, nursing, and fecal bacteria. Non-breeding adults were housed in single sex groups of ca. 3 individuals (representing the same line type) per cage (open type), with bedding and an upturned coconut shell provided for shelter and enrichment. Rodent pellets (standard rodent chow: energy: 12.7MJ, 24% protein, 3% fat, 4% fiber; Labofeed H, Kcynia, Poland) and water were available *ad libitum*. Animals from both selection treatments were maintained in the same rooms, and thus were exposed to shared environmental bacteria under standard maintenance conditions.

### Study design

In spring 2020, 224 non-breeding adult bank voles were sampled from the breeding colony and housed singly in individually ventilated cages for the duration of the experiment. The voles were randomly allocated to diet treatments (standard / energy restricted), with 7 individuals of each sex (male / female) from each of eight selection lines (four A lines: A1-A4, and four C lines: C1-C4) in each diet treatment. The animals were distributed randomly with respect to the cage position in the ventilation system.

The study design (Fig. 1) entailed three phases. In **phase 1, baseline**, days 1-6 of the experiment, all individuals were fed the standard diet used in the colony while being acclimated to the individually ventilated cages and we assessed the initial vole physiology and fecal microbiota. In **phase 2, diet treatment,** days 7-33, animals were divided among two diet treatments: (1) standard diet (SD) or (2) dietary restriction through cellulose dilution of the diet (CR). The SD treatment continued to receive the same rodent pellets that all voles received in phase 1. The CR group was acclimated for six days with a 15% cellulose-diluted diet, to which the manufacturer added 15% of cellulose (hereafter: CR15), and then, for 21 days, provided with a 30% cellulose-diluted diet (hereafter: CR30). Finally, all animals continued to **phase 3, recovery period**, days 34-47, during they were fed a standard diet again. The metabolizable energy content of the diets in kJ/g dry mass was 14.37 for SD, 12.12 for CR15, and 9.86 for CR30.

On the last day of each phase (P), we collected phenotypic measurements from the bank voles and sampled feces for microbiota analyses. Hereafter, the measurements and samples obtained at these three time points are referred to as P1 (baseline, day 7), P2 (end of diet treatment, day 34), and P3 (end of recovery, day 47) (Fig. 1).

### Sampling and measurement techniques

**Daily food consumption** and calorie intake were regularly approximated by provision of a weighed quantity of food pellets, weighing the remaining food after 3-4 days and averaging for daily consumption. **Body mass** was measured using a digital scale to 0.1g accuracy.

**Resting metabolic rate (RMR)** was measured at room temperature as previously (Sadowska et al. 2015a; Hseiky et al. 2025) by placing animals individually into respirometric chambers (850 ml) without food or water and measuring the rate of oxygen consumption (VO_2_; ml O₂ min⁻¹) using an open-flow, positive-pressure respirometric system (see SI for details). Outflow was sampled every 13 minutes and analyzed using a CO_2_ analyzer (LI-850, LI-COR Biosciences, Lincoln, NE, USA) and an oxygen analyzer (AMETEK S3AII, Applied Electrochemistry, Pittsburgh, PA, USA) for a maximum of 17 cycles after excluding the first two cycles. RMR was operationally defined as the mean value of the three lowest VO_2_ values recorded for a given individual. Movement activity of the animals were evaluated with MAD-1 gravimetric detectors (Sable Systems, Las Vegas, NV, USA).

**Fecal samples** were collected for microbiota analyses from an empty, clean cage and samples were stored at -80 ֯C until DNA extraction.

On the day after the end of P3 (day 48), bank voles, euthanized with isoflurane inhalation overdose, were dissected, and **digestive tract mass** (stomach, caecum, and intestine) was measured. The remaining carcass (excluding the digestive tract) was dried to constant mass and subjected to lipid extraction using a hot Soxhlet apparatus to quantify total **body fat**, calculated as the difference between dry carcass mass before and after fat extraction.

### DNA extraction and sequencing

We sequenced 480 fecal samples from the 169 vole individuals that had the most complete data on variables of interest throughout the experiment (P1-P3), balancing treatment groups, sexes, and selection lines. Total genomic DNA was extracted from the fecal samples using DNeasy Power Soil Pro kit (Qiagen, Germany) using the manufacturer’s instructions. Additionally, 12 negative controls were collected in the isolation and library preparation steps. We prepared amplicon sequencing libraries for the V4 region of the 16S rRNA gene following a two-step PCR protocol. In the first step, we amplified 16S rRNA variants using primers 515F and 806R with Illumina adapter tails. In the second PCR, we completed adapters and added unique indexes. The quality-verified samples were pooled and quality checked using BioAnalyzer 2100 (Agilent), and submitted to the Swedish National Genomics Infrastructure for paired-end sequencing on Illumina NovaSeq SPrime flow cells (2 x 250 bp read mode) as in (Nowak et al. 2025).

### Read data processing

Demultiplexed forward and reverse reads had primers verified and then trimmed using a custom script, and were then assembled into contigs using PEAR (Stamatakis et al. 2014). Further analyses were performed in QIIME2 v. 2023.5.1 (Bolyen et al. 2019). After quality filtering, denoising, chimera removal, and trimming, the feature table consisted of 18,974,257 sequences with 4,012 amplicon sequence variants (ASVs). Eight extraction blanks and one vole sample failed sequence-quality criteria and were excluded from downstream analyses, including contaminant assessment. Taxonomy was assigned to the ASVs using a naïve Bayes classifier (QIIME2 sklearn, default parameters) pretrained on the SILVA v. 138_99 16S rRNA database. A midpoint rooted tree was assembled using align-to-tree-mafft-fasttree plugin. After taxonomy assignment, the feature table was cleaned from eukaryotes, Archaea, chloroplasts, mitochondria, and unassigned phyla.

Contaminant ASVs (n=29) were identified and excluded from the feature table using the frequency method in package decontam v3.16 (Davis et al. 2018) in program R (version 4.2.2 (R Core Team 2018)). Data were rarefied to 10,058 reads per sample and this normalized feature table was used for further analyses except where stated. During rarefaction, 4 samples were dropped, leaving 474 samples (from 168 voles; N= 158, 159, and 157 samples from phases P1, P2, and P3, respectively) with 4,767,492 (25.91%) total reads and 3,960 ASVs.

**Alpha diversity** of the gut microbiota was estimated by calculating observed number of ASVs and Shannon index for each sample. Pairwise sample dissimilarities were calculated using three different **beta diversity** metrics: unweighted UniFrac distance (community membership), and Bray–Curtis dissimilarity and weighted UniFrac distance (community structure).

### Statistical analyses

Statistical analyses were performed using R v. 4.2.2 (R Core Team 2018) and v. 4.4.2 (R Core Team 2024).

### Host response to temporary dietary restriction

Overall differences in body mass, food consumption, and resting metabolic rate (RMR) between selection line types (C/A) at baseline were initially compared using Welch two sample t-tests.

To test whether dietary restriction influenced the voles differently depending on the selection line type, we estimated linear mixed effects models using nlme v. 3.1.166 (Pinheiro et al. 2024; Pinheiro and Bates 2000)). This allowed us to account for structures in the data owing to replicate selection lines and longitudinal sampling from the same individuals. A separate model was built for RMR, body mass, and food consumption. All response variables were z-transformed for comparability, and back-transformed for plotting. The sample sizes varied among models due to some cases of missing data (due to data not recorded, uncertain vole identity, unreliable or biologically impossible measurement result), and are listed separately for each analysis in SI. All models included the fixed effects of the dietary treatment (SD/CR), selection line type (C/A), and experiment phase (P1/P2/P3), and all their two- and three-way interactions. As additional control variables we included sex (female, male) in all the models, body mass (measured in the same phase) in the models for food consumption and RMR, and activity for the RMR model. Random effects in all the above models included the random intercept of vole individual nested within selection line (C1-4, A1-4) to account for repeated measurements.

The cross-sectional fat mass and total digestive organ mass were modeled separately using lme4 v.1.1.36 (Bates et al. 2015), with the fixed terms selection, diet treatment, their interaction, sex, carcass dry mass, and dissection delay (see SI), with replicate line included as a random intercept.

### Effects of dietary restriction on gut microbiota

Bacterial alpha diversity was examined with the same general model structures as detailed above for the host physiological traits, but the count of observed ASVs was modeled using glmmTMB v. 1.1.10 (Brooks et al. 2017) using a negative binomial error distribution, while Shannon index was modeled using a gaussian error distribution in lme4 v. 1.1.36 (Bates et al. 2015).

To quantify longitudinal (within-individual) changes in community composition, we used the q2-longitudinal plugin (Bokulich et al. 2018) in QIIME2 to compute the **first distances**, i.e. the magnitude of change in beta diversity among successive time points. We computed the first distances with random replicate handling from baseline P1 to P2 (effect of diet treatment on microbiota), and from P1 to P3 (completeness of recovery) for each of the three different beta diversity metrics: unweighted and weighted UniFrac distance and Bray–Curtis dissimilarity. The first distances were used as response variables in linear mixed models in lme4. We computed two separate models for A) the distance from P1 to P2, and B) the distance from P1 to P3. Thus, for model A, larger distances in the CR treatment compared to the SD treatment would indicate a longitudinal shift in microbiota community due to dietary restriction. For model B, distances close to zero would indicate complete community recovery to near baseline. Both models included treatment × selection interaction and main effects, and sex. The random intercept of line was included as a random effect; when it had a zero variance and caused non-convergence, a linear model with the same fixed terms was estimated instead (likelihood ratio test of nested models with and without random effect of line, all Χ^2^=0, P=1).

To identify the taxa that differ in abundance among treatment groups, we used ancombc2 (ANCOMBC v. 2.8.1 (Lin and Peddada 2020, 2024; Lin et al. 2022) with default settings and structural zeros detected based on diet treatment. We estimated differential abundances separately for phases P2 and P3 on genus and family level, with each model constructed with the fixed terms treatment × selection interaction and main effects, and sex. The random formula was set to NULL due to nonconvergence resulting from singularities. Additionally, we examined the main effect of diet treatment on differentially expressed families at P2 to describe the overall effects of the diet treatment. Bacterial taxa that failed sensitivity analyses for the variable of interest (treatment × selection or treatment) were excluded from the results. Taxa identified as differentially abundant for the interaction term were examined by modeling the log10-transformed abundance (+1e-6 to account for zeros) of the family/genus within a sample as a function of treatment × selection and sex, and the random intercept of line with a gaussian error distribution in lme4 v. 1.1.36 (Bates et al. 2015).

For the final models, we estimated marginal means using emmeans (joint tests) v.1.10.7 (Lenth 2025) and plotted the results for the diet treatment × selection (× phase) interaction with ggplot2 v.4.0.1 (Wickham 2016). Full modeling results are provided in SI.

## Results

Overall, we found limited support for our general hypothesis that a higher selected metabolic rate should influence the magnitude of changes in host physiology and gut microbiota under temporary dietary restriction, summarized in Table 1.

**Table 1.**
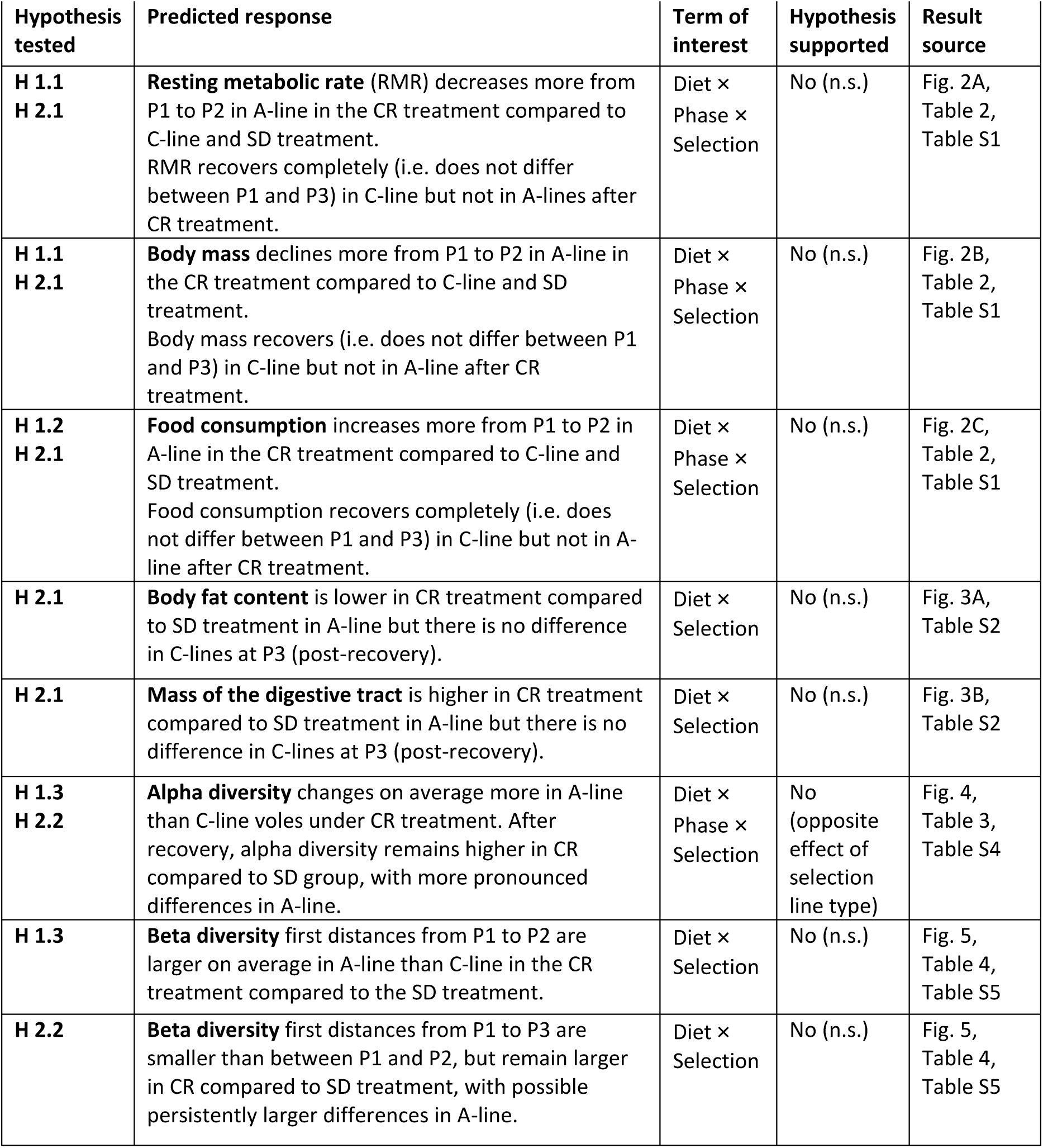
Summary of hypotheses, predicted responses, and outcomes. Overview of statistical tests evaluating the primary hypothesis> whether bank voles from high-metabolism selection lines (A) differed from control lines (C) in their physiological and gut microbiota responses to dietary restriction (CR) relative to the standard diet (SD). Experimental phases were P1 (baseline), P2 (end of dietary restriction), and P3 (end of recovery). “Term of interest” indicates the model term used to evaluate each prediction. Support for each hypothesis is evaluated based on statistical significance of the term of interest. N.s. = non-significant (P>0.05). Specific hypotheses are referred to by number corresponding to the hypotheses proposed in the introduction. Please refer to the Figure 1 for experimental design and methods for detailed description.

### Host response to temporary dietary restriction

#### Effects of metabolic rate selection in baseline values

In line with earlier studies, bank voles selected for a fast metabolism (A-line voles) had a 8% higher body mass at baseline than control (C-line) voles (mean body mass at baseline (P1) for A: 23.3 g, C: 21.5 g, t = -3.174, df = 195.7, P = 0.002), their apparent food consumption was on average 39% higher (mean of food consumed per day without body mass correction for A: 5.9 g (84 kcal), C: 4.2 g (61 kcal), t = -12.542, df = 217.2, P < 0.001), and their resting metabolic rate was 20% higher than that of C-line voles (mean VO_2_ for A: 1.71 ml O_2_/min, C: 1.37; t = -8.884, df = 185.38, P <0.001; Fig. S2).

#### Effects of diet treatment on food and calorie intake

The effectiveness of the CR diet treatment was confirmed by comparing food and calorie intake at the end of the dietary restriction at P2. CR treatment increased food intake but still caused an apparent calorie intake reduction relative to SD in both selection line types. At P2, the average daily food intake of A-line voles was ca. 13% higher but calorie intake 21% lower on CR diet than on SD diet (SD: 77±14 kcal/day, CR: 61 ± 9 kcal), whereas in C-line voles daily food intake increased on average 8 % while calorie intake decreased by 23 % on CR diet (SD: 60±17 kcal, CR: 46 ± 9 kcal; Fig. S2).

#### Effects of diet treatment on host characteristics

##### Physiological traits and energy consumption

Contrary to our primary hypothesis, the effect of the diet treatment did not differ for the selection directions in terms of body mass, food consumption, or resting metabolic rate (non-significant three-way interaction of selection, treatment, and phase for each, Table 2). A reduction in RMR was seen in A-line voles of both treatments from baseline to subsequent phases, perhaps indicating an acclimation response to the measurement situation, while the RMR of C-line voles remained similar across the phases and treatments (Fig. 2A, Table S1). RMR was overall lower in CR treatment than in SD-treatment in A-lines at P2, with some residual effect at P3, while RMR differed little between treatments in C-line voles. Body mass declined (Fig. 2B) and food consumption increased (Fig. 2C) in both selection directions from P1 to P2 and returned to near baseline by P3 (two-way interactions of phase with selection and diet treatment (Table 2; Table S1).

**Figure 2.**
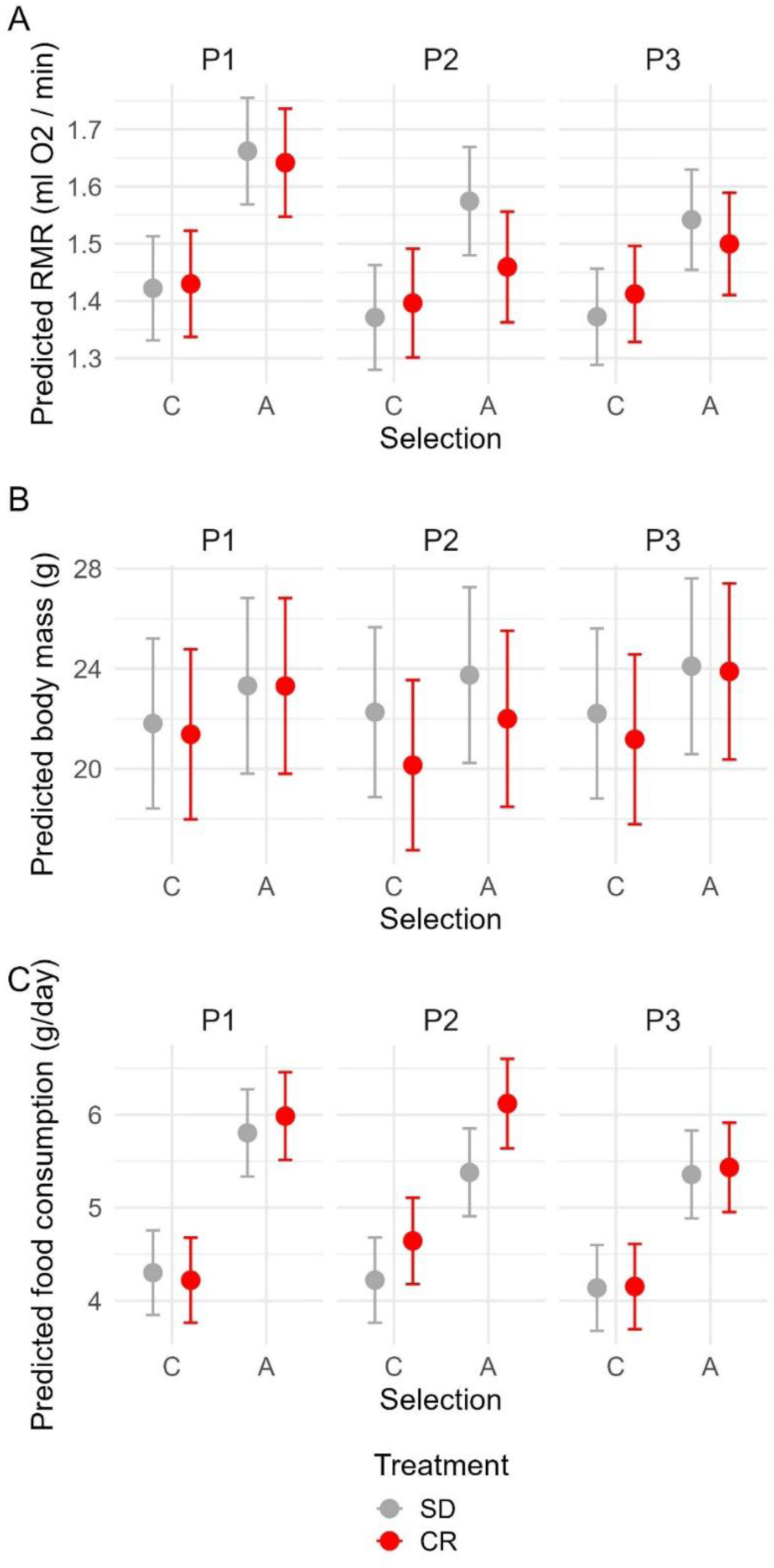
Effects of selection history, diet treatment, and experimental phase on host physiological traits. Model-predicted, back-transformed marginal means for (A) resting metabolic rate (RMR), (B) body mass, and (C) daily food consumption across experimental phases (P1-P3), diet treatments (SD and CR), and selection line types (A and C). Points represent estimated marginal means from mixed-effects models and error bars indicate 95% confidence intervals.

**Table 2.** Joint tests of fixed effects for host physiological traits. Results of mixed-effects models examining effects of diet treatment (SD vs. CR), experimental phase (P1-P3), selection line type (A vs. C), and their interactions on resting metabolic rate (RMR), body mass (BM), and food consumption (FC). Shown are numerator and denominator degrees of freedom (df), F-ratios, and associated P-values from joint tests of estimated marginal means. All models were estimated with compound symmetry residual covariance structure. A separate residual variance was estimated for each phase in RMR and FC models, while a homogeneous residual variance structure was used in the BM model. Full model outputs are provided in Supplementary Table S1.

| Model term | Resting metabolic rate |  |  |  | Body mass |  |  |  | Food consumption |  |  |  |
| --- | --- | --- | --- | --- | --- | --- | --- | --- | --- | --- | --- | --- |
|  | df1 | df2 | F-ratio | P-value | df1 | df2 | F-ratio | P-value | df1 | df2 | F-ratio | P-value |
| Treatment | 1 | 210 | 0.838 | 0.361 | 1 | 210 | 5.890 | 0.016 | 1 | 209 | 4.473 | 0.036 |
| Phase | 2 | 374 | 23.658 | **<0.001** | 2 | 401 | 24.449 | **<0.001** | 2 | 405 | 19.161 | **<0.001** |
| Selection | 1 | 6 | 13.192 | **0.011** | 1 | 6 | 0.911 | 0.377 | 1 | 6 | 35.704 | **0.001** |
| Treatment:Phase | 2 | 374 | 1.353 | 0.260 | 2 | 401 | 30.613 | **<0.001** | 2 | 405 | 13.686 | **<0.001** |
| Treatment:Selection | 1 | 210 | 4.830 | **0.029** | 1 | 210 | 0.500 | 0.480 | 1 | 209 | 1.010 | 0.316 |
| Phase:Selection | 2 | 374 | 7.404 | **0.001** | 2 | 401 | 4.680 | **0.010** | 2 | 405 | 6.340 | **0.002** |
| Treatment:Phase:Selection | 2 | 374 | 1.758 | 0.174 | 2 | 401 | 0.568 | 0.567 | 2 | 405 | 0.655 | 0.520 |

##### Body fat content

A further indication of a full physiological recovery from dietary treatment is the body fat content, which was similar in voles from the two diet treatments at the end of the recovery period (treatment:selection (Χ^2^ =1.563, P=0.211) (Fig. 3A, Table S2). A-line voles had on average a 16% lower relative fat mass compared to the C-line voles in both treatments (P=0.039; Fig. 3A, Table S2).

**Figure 3.**
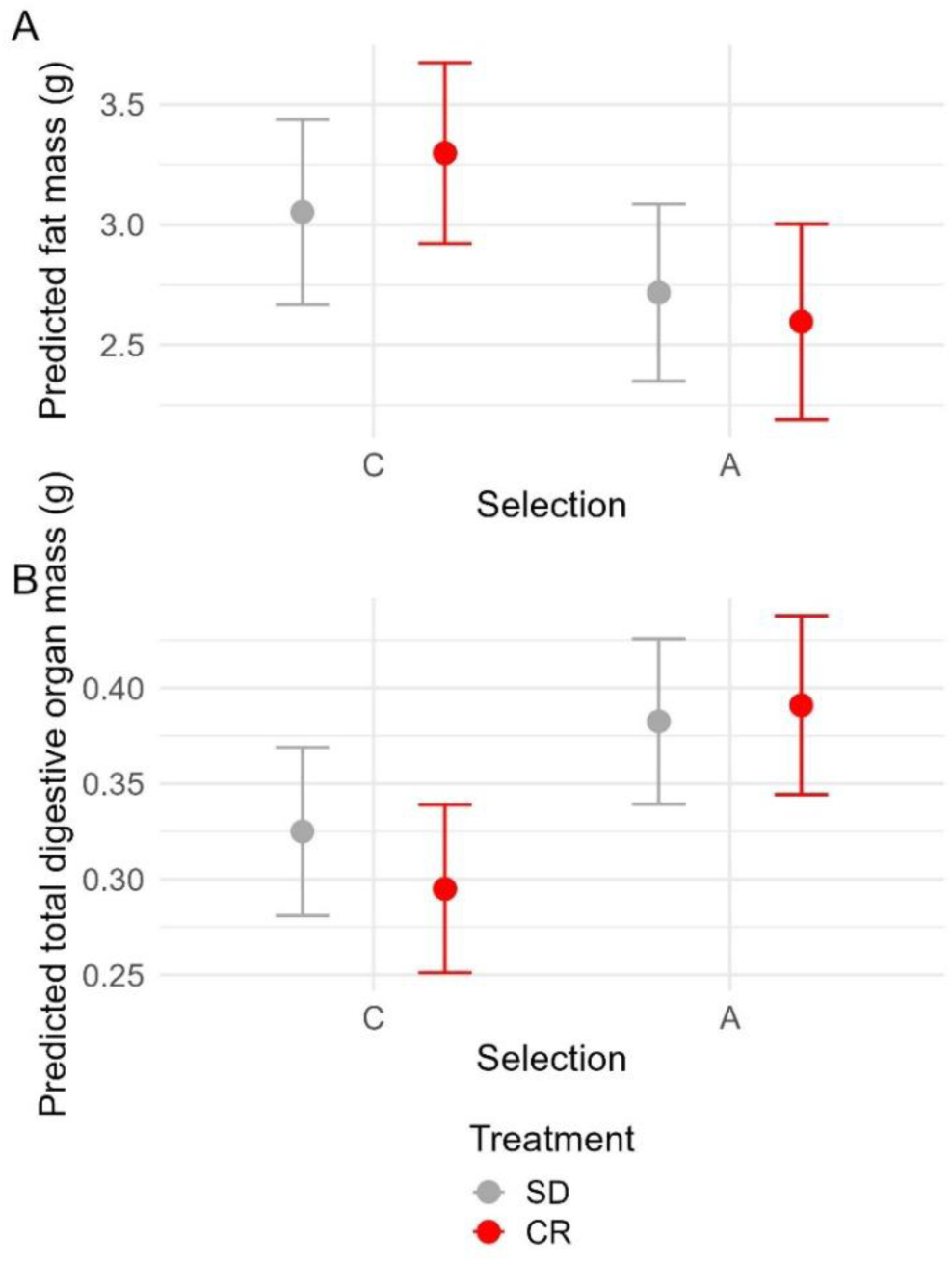
Diet treatment × selection effects on body fat and digestive tract mass at the end of recovery. Model-predicted marginal means for (A) body fat mass and (B) digestive tract dry mass in high-metabolism (A) and control (C) bank voles at the end of the recovery period (P3) following standard diet (SD) or dietary restriction (CR) treatment. Points show estimated marginal means and error bars indicate 95% confidence intervals. The diet treatment × selection interaction was not significant for either trait.

##### Digestive tract mass

There was no evidence that A-line and C-line voles differed in their digestive tract dry mass response to dietary restriction (treatment:selection interaction effect on combined organ mass: Χ^2^ =2.329, P=0.127; Fig. 3B; Table S2). A-line voles had on average 20% heavier digestive tract (C-lines: 0.310 g vs. A-lines: 0.387 g, P= 0.023, Table S2). Similar trends were found for the selection effect when looking at the stomach, caecum, and intestines separately (Fig. S3).

### Effects of dietary restriction on gut microbiota

#### Bacterial community of the bank voles

The gut microbiota of bank voles across all groups comprised 12 phyla (Fig. S4), which were dominated by the *Bacteroidota* (=*Bacteroidetes*) and *Bacillota* (=*Firmicutes*) (average proportions: 0.47 and 0.43, respectively) followed by *Desulfobacterota* (0.02) and *Campilobacterota* (0.02). The most abundant families were *Muribaculaceae* (0.40), *Lactobacillaceae* (0.17), and *Lachnospiraceae* (0.11) (Table S3).

#### Alpha diversity

Bacterial alpha diversity increased in response to the cellulose-diluted diet in both selection line types and decreased after the recovery period to near baseline levels. For both observed number of ASVs and Shannon index, the treatment effect differed significantly depending on the selection line type (significant three-way interaction of selection:treatment:phase Table 3; Table S4). Contrary to our hypothesis, however, the initial response (P1-P2) was significantly larger for the C-line compared with the A-line voles, and diversity remained slightly higher in the C-line compared with the A-line voles after the recovery period (P3) (Fig. 4, Table 3). Notably, the treatment effect at the end of the diet treatment (P2) was minor with regard to observed ASVs (pairwise comparison of marginal means at P2 for SD vs. CR for both C-line and A-line voles: P=1; Fig. 4A) compared with a large difference in Shannon index between treatments (pairwise comparison of marginal means at P2 for SD vs. CR for C-line voles P<0.001, A-line voles P=0.050; Fig. 4B), indicating that the diet treatment primarily affects community dynamics (relative abundances of species), with community membership (gain/loss of species) playing a small role.

**Figure 4.**
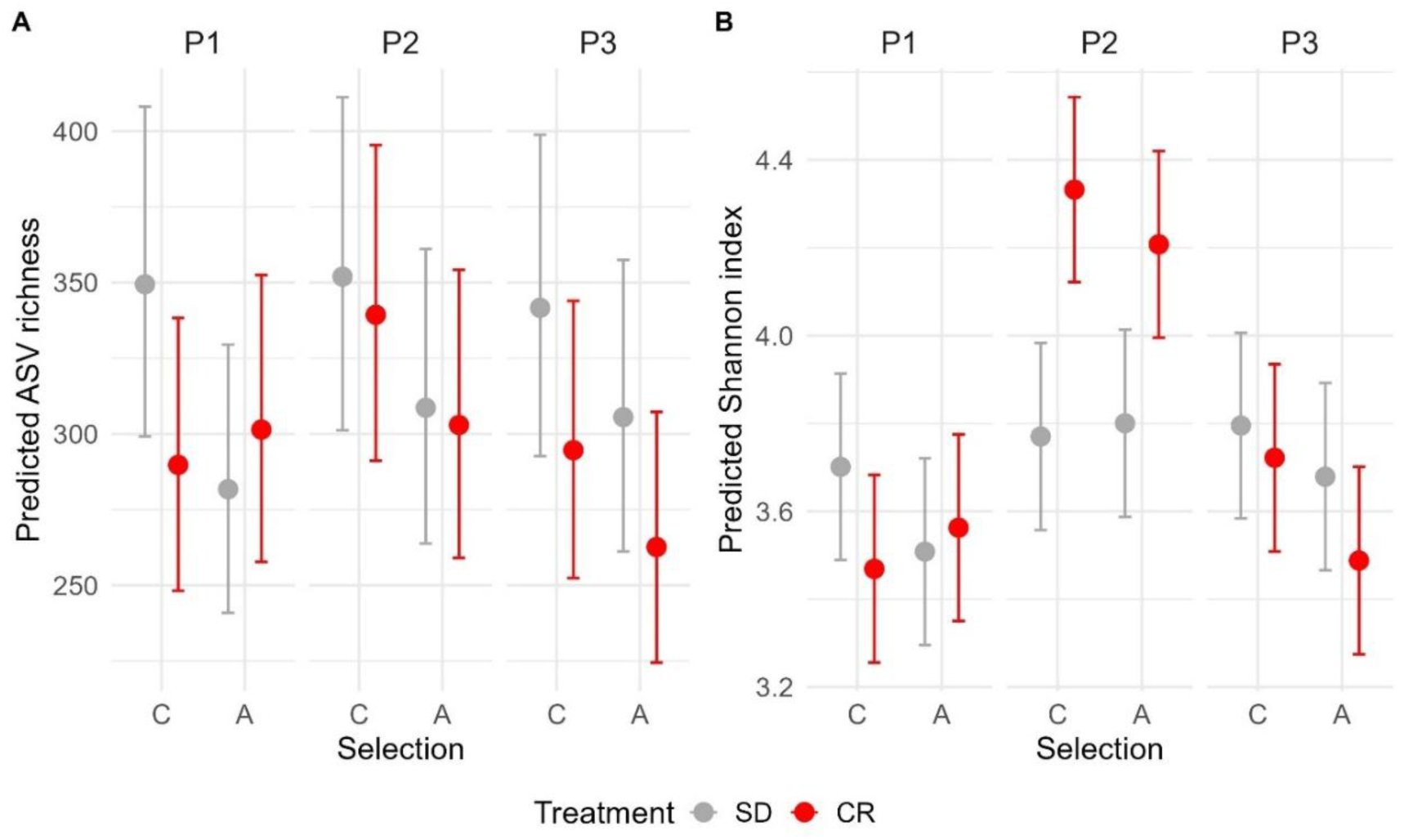
Effects of dietary restriction and recovery on gut bacterial alpha diversity. Estimated marginal means (±95% confidence intervals) from mixed-effects models showing changes in bacterial alpha diversity across selection line types (A and C), diet treatments (SD and CR), and experimental phases (P1-P3) in (A) Observed amplicon sequence variant (ASV) richness and (B) Shannon diversity index. Points indicate model-predicted means and error bars denote 95% confidence intervals. Estimates are adjusted for sex and random effect of vole individuals nested within replicate lines.

**Table 3.** Effects of dietary restriction, selection history, and phase on gut bacterial alpha diversity. Results of longitudinal models testing variation in bacterial alpha diversity, measured as Observed ASV richness and Shannon index, across experimental phases (P1-P3), diet treatments (SD vs. CR), and selection line types (A vs. C). Full model results are presented in Supplementary Table S4.

| Model term | Observed ASVs |  |  |  | Shannon index |  |  |  |
| --- | --- | --- | --- | --- | --- | --- | --- | --- |
|  | df1 | F-ratio | X<sup>2</sup> | P-value | df1 | df2 | F-ratio | P-value |
| Selection | 1 | 1.703 | 1.703 | 0.192 | 1 | 6.00 | 0.703 | 0.434 |
| Treatment | 1 | 1.472 | 1.472 | 0.225 | 1 | 155.60 | 1.888 | 0.171 |
| Phase | 2 | 6.382 | 12.764 | **0.002** | 2 | 303.95 | 67.705 | **<0.001** |
| Selection:Treatment | 1 | 0.475 | 0.475 | 0.491 | 1 | 155.67 | 0.001 | 0.970 |
| Selection:Phase | 2 | 0.277 | 0.554 | 0.758 | 2 | 303.95 | 1.494 | 0.226 |
| Treatment:Phase | 2 | 3.404 | 6.808 | **0.033** | 2 | 303.92 | 33.513 | **<0.001** |
| Selection:Treatment:Phase | 2 | 4.420 | 8.840 | **0.012** | 2 | 303.95 | 4.231 | **0.015** |

By chance, alpha diversity was lower at baseline in the C-line bank voles assigned to CR treatment compared to SD treatment (Fig. 4), but this difference was not statistically significant (pairwise comparison of marginal means for SD vs. CR for C-line voles at P1 for observed ASVs and Shannon index: both P>0.6).

#### Beta diversity

Dietary restriction caused a longitudinal change in the bank vole gut bacterial community composition in both selection line types (Fig. 5, Table 4, Fig. S5). The change from a standard diet to a cellulose-diluted diet (P1 to P2) caused a significant change in the gut microbial community compared to the group staying on standard diet (effect of diet treatment: P≤0.009 for first distances in all beta diversity metrics; Table 4). There was no statistically significant effect of selection line or sex on the change in gut microbiota composition (all P>0.07, Table S5).

**Figure 5.**
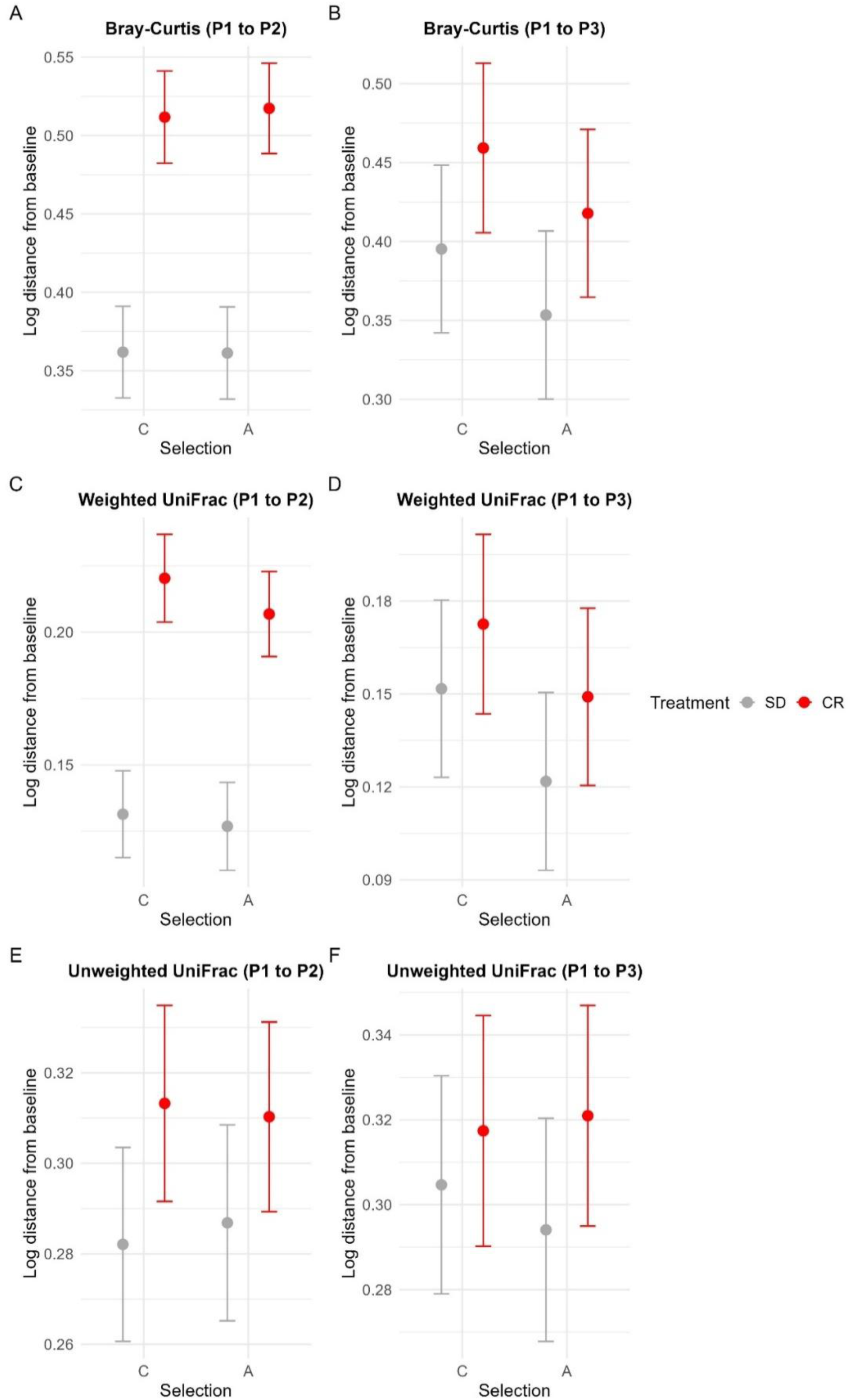
Longitudinal changes in gut microbiota composition during dietary restriction and recovery. Model-estimated first distances quantifying within-individual changes in gut bacterial community composition relative to baseline (P1). Panels show Bray-Curtis dissimilarity (A, B), weighted UniFrac distance (C, D), and unweighted UniFrac distance (E, F) between P1 and the end of dietary restriction (P2; A, C, E) or the end of recovery (P3; B, D, F). Results are shown separately for control (C) and high-metabolism (A) selection lines under standard diet (SD) and dietary restriction (CR). Points represent estimated marginal means and error bars indicate 95% confidence intervals. Larger first distances indicate greater divergence from baseline microbiota composition.

**Table 4.** Effects of dietary restriction on longitudinal gut microbiota compositional change. Results of models evaluating first distances (log-transformed) for Bray-Curtis, weighted UniFrac, and unweighted UniFrac. The P1-P2 comparison quantifies microbiota change from baseline to the end of dietary restriction. The P1-P3 comparison reflects the completeness of recovery as remaining difference from baseline after the recovery period. Effects of selection line type (A vs. C), diet treatment (SD vs. CR), and their interaction were modeled. Full model outputs are provided in Supplementary Table S5.

|  | P1-P2 first distance |  |  |  | P1-P3 first distance |  |  |  |
| --- | --- | --- | --- | --- | --- | --- | --- | --- |
|  | df1 | df2 | F-ratio | P-value | df1 | df2 | F-ratio | P-value |
| **Bray-Curtis** |  |  |  |  |  |  |  |  |
| Selection | 1 | 5.98 | 0.030 | 0.880 | 1 | 6.00 | 1.930 | 0.210 |
| Treatment | 1 | 141.21 | 172.540 | **<0.001** | 1 | 136.12 | 17.560 | **<0.001** |
| Selection:Treatment | 1 | 141.21 | 0.070 | 0.790 | 1 | 136.13 | 0.000 | 0.990 |
| **Unweighted Unifrac** |  |  |  |  |  |  |  |  |
| Selection | 1 | 147.00 | 0.008 | 0.930 | 1 | 143.00 | 0.077 | 0.782 |
| Treatment | 1 | 147.00 | 6.962 | **0.009** | 1 | 143.00 | 2.428 | 0.121 |
| Selection:Treatment | 1 | 147.00 | 0.139 | 0.710 | 1 | 143.00 | 0.312 | 0.578 |
| **Weighted Unifrac** |  |  |  |  |  |  |  |  |
| Selection | 1 | 147.00 | 1.305 | 0.255 | 1 | 6.00 | 2.798 | 0.145 |
| Treatment | 1 | 147.00 | 114.141 | **<0.001** | 1 | 136.15 | 7.214 | **0.008** |
| Selection:Treatment | 1 | 147.00 | 0.318 | 0.574 | 1 | 136.15 | 0.129 | 0.720 |

The diet-induced shift in microbiota communities was largely reversed when the standard diet was resumed (Fig. S5). Some residual changes at individual level remained in the relative abundances of taxa, as indicated by P1 to P3 first distances in the weighted metrics (Bray-Curtis distances, Weighted UniFrac P ≤ 0.008; Table 4, Fig. 5; Table S5). In these, the change from baseline to post-recovery communities remains significantly larger in the dietary restriction compared to the standard diet treatment. In contrast, no differences were found among diet treatments in Unweighted UniFrac first distances (P>0.1; Table 4, Fig. 5), indicating that the taxonomic assembly was fully reversed back to baseline. No significant effect of selection line or sex was detected in the recovery (P1 to P3) of gut microbiota community composition.

#### Effects of dietary restriction on community composition among selection line types

Differential abundance analyses (ANCOMBC2) indicated overall reversible changes in microbiota in response to the diet treatment. The treatment effect was largely similar for the selection lines, with the exception of one family *Helicobacteraceae* (mean relative abundance: 0.019) and genus *Helicobacter* within it. At the end of the diet treatment (P2), these taxa were enriched ca. 6.6 fold in the CR treatment compared to SD in the C-line voles, while this effect was absent in the A-line voles (treatment:selection for *Helicobacteraceae* logfold change LFC=-1.751, SE=0.489, P_fdr_ = 0.024; *Helicobacter* LFC=-1.751, SE=0.485, P_fdr_ = 0.040). However, the baseline levels of these taxa were higher in A-line voles (ca. 3.9-fold higher relative abundance in A- than C-lines in SD). This interaction was further confirmed with a mixed effects model of the relative abundance of *Helicobacteraceae* (treatment:selection P=0.004; Fig. 6, Table S6). By P3, treatment:selection interaction influenced no taxa significantly.

**Figure 6.**
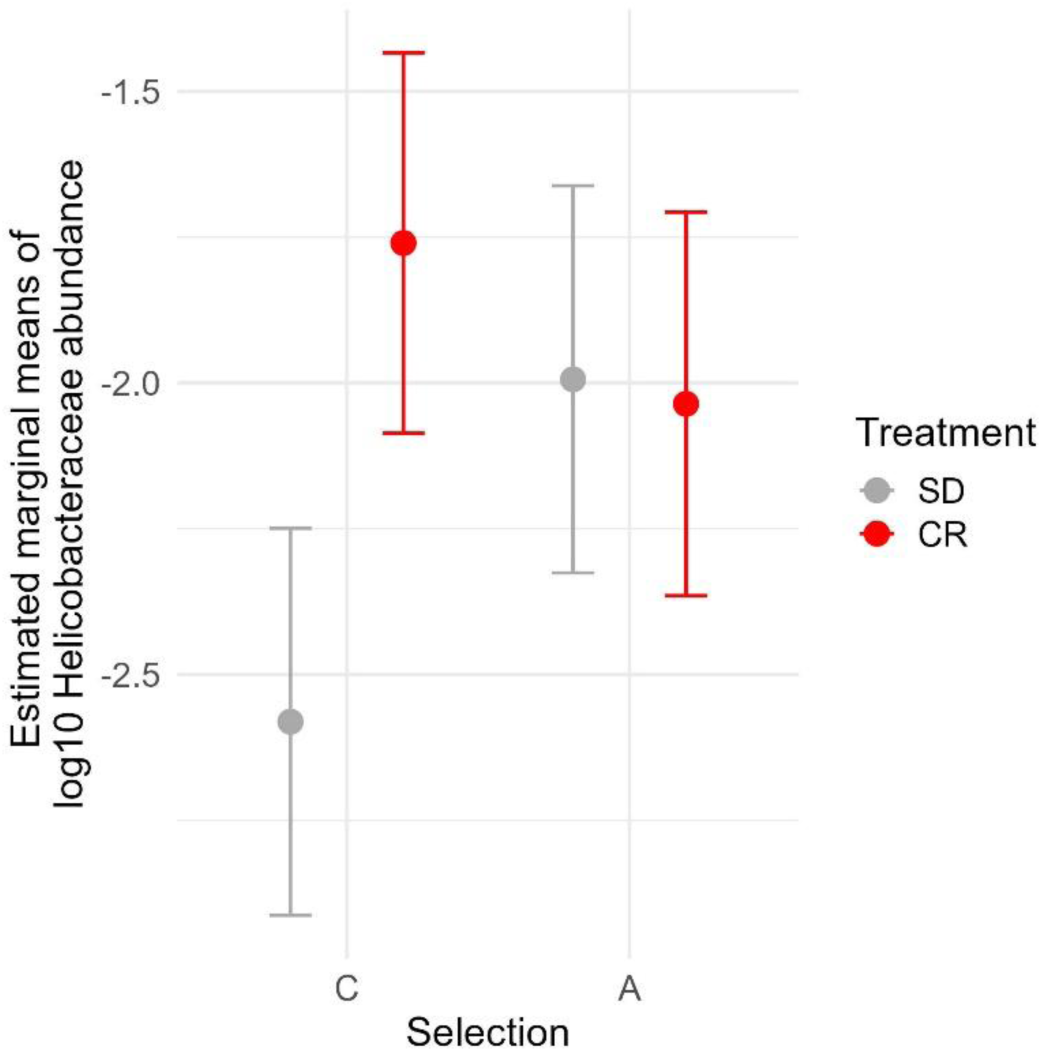
Response of *Helicobacteraceae* to dietary restriction modified by selection. Estimated marginal means (±95% confidence intervals) of the log10-transformed relative abundance of *Helicobacteraceae* (log10(abundance + 1 × 10⁻⁶)) at the end of the dietary restriction period (P2) in bank voles from control (C) and high-metabolism (A) selection lines exposed to either standard diet (SD) or dietary restriction treatment (CR). Points represent model-predicted means and error bars indicate 95% confidence intervals.

The diet treatment altered the abundances of a number of taxa independent of selection line type (Fig. S6, Table S7). Of the most abundant families, *Lachnospiraceae* was significantly enriched in the CR treatment compared to the SD treatment at P2 (log fold change LFC= 0.549, P_fdr_<0.001). Another two rare families (≤0.003) were also enriched in CR. Simultaneously, ten families were reduced, including five relatively common taxa (relative abundance 0.01-0.17; LFC < -1.9; P_fdr_ <0.001): *Christensenellaceae*, *Bifidobacteriaceae*, *Erysipelotrichaceae, Lactobacillaceae, and Spirochaetaceae* (Fig. S6). Out of these, *Christensenellaceae* was still reduced in the CR treatment group after recovery at P3 (LFC= -2.541, SE=0.337, W=-7.538, P_fdr_<0.001).

## DISCUSSION

Dietary restriction influences both host physiology and microbiota composition and function, but the extent to which these effects are modulated by intrinsic host energy needs is poorly understood. In this study, we tested experimentally whether a higher inherent energy need of a host (A-line bank voles artificially selected for a fast metabolism vs. C-line controls) magnifies the effects of temporary dietary calorie reduction (dilution of diet with 30% cellulose fiber) on host physiology and gut microbiota. We observed an expected impact of dietary restriction on most of the studied host traits and gut microbiota. However, we found no support for our hypothesis that animals with a higher intrinsic energy demand (A-line voles) should respond more strongly than control (C-line) voles to reduced energy availability. Indeed, A-line microbiota appeared more resistant to dietary change. Overall, the results indicate that the influence of host genetics on microbiota composition and function under nutritional stress is limited and surpassed by environmental factors.

Energy availability in the environment has been suggested to influence the competitive ability and physiological response of organisms with different energy demands (Hall et al. 1992; Brandl et al. 2023). Considering this, our findings of only minor differences between the selection lines in the strength of physiological and morphological responses (body mass, food consumption, RMR, digestive tract mass) to dietary restriction are surprising. They are also in contrast with an earlier laboratory study, in which an inherited difference in metabolic rate significantly influenced the response to dietary restriction in mice (Brzęk et al. 2012). In these mice, dietary restriction with 30% reduced food quantity reduced basal metabolic rate more in mouse lines selected for high basal metabolic rate compared with lines selected for low basal metabolic rate. Notably, (Brzęk et al. 2012) quantified direct change in the selected trait, while we examined coevolved metabolic and other traits rather than the trait under directional selection (V_O2, swim_), which may partially explain the difference in effect size. In our study, the bank voles were also able to partially compensate for the reduction in digestible calories by increasing their food intake and potential adjustments in the microbiota to derive more energy from the food. It is plausible that such a concerted response of the host behavior and microbiota can overcome genetic differences in energy needs.

The gut microbiota could compensate for the reduction in readily digestible energy by changes in community structure and metabolism that allow more efficient extraction of energy from food, e.g. via increased production of short chain fatty acids (den Besten et al. 2013). If this compensation was more pronounced for the A-line relative to C-line voles, that might explain the apparently similar physiological response of the selection lines to the diet treatment. Instead of a stronger response in the A-line voles’ microbiota, however, we observed an opposite trend: greater changes in alpha diversity were apparent in the C-line voles compared to A-line voles at P2, suggesting a restructuring of the microbial community. This change was insufficient to be manifest as a significant change in community composition among selection line types. The apparently subdued response of the A-line gut microbiota may indicate an effect of selection on the resilience to a reduction in diet quality, or the control of the host physiology on the gut microbiota. In a previous study, we similarly observed stronger changes in the C-line bank voles’ gut microbiota when the lines were released into semi-natural field conditions (Hanhimäki et al. 2022), while in standard rearing conditions, the lines have similar gut microbiota. Taken together, our findings indicate higher resistance (reduced flexibility) of A-line voles’ gut microbiota to disturbance, while the resilience of the microbiota (ability to return to the original state after disturbance) appears similar for the selection lines. Host evolutionary background thus influenced specifically the microbiota flexibility in subtle but repeatable ways.

Microbiota can respond to environmental change faster than the host as microbial communities rapidly restructure in response to changing conditions (Voolstra and Ziegler 2020). By mediating interactions between the host and its environment, this microbiota flexibility may buffer stress and facilitate host phenotypic plasticity (Lynch and Hsiao 2019). Here, we observed substantial reversible plasticity in both host physiology and gut microbiota: body mass, food consumption, RMR, and microbiota composition returned close to baseline within two weeks after dietary restriction ended. The increase in Shannon diversity with only minor changes in ASV richness suggests that dietary restriction primarily altered the relative abundances of bacterial taxa rather than community membership. Although previous studies have shown that gut microbiota can shift rapidly following dietary change and recover after return to the original diet (E. M. Anderson et al. 2022; Carmody et al. 2015), recovery from significant stress may remain incomplete (Huang et al. 2022; Spor et al. 2011). The near-complete recovery of the gut microbiota observed here indicates highly flexible community dynamics in bank voles exposed to an energetic challenge.

Given that there was no marked difference between the selection lines in the effect of the dietary restriction despite differences in energy demand, and the microbiota of A-line voles demonstrated no obvious compensatory changes in A-lines, the mechanisms underlying the apparently proportionately higher resistance of the A-line voles to dietary restriction remains unclear. Possible explanations include altered host physiological / unmeasured metabolic processes, behavioral adaptation (e.g. reduced activity), and functional shifts in the gut bacteria. The microbiota has coevolved with the host lines in a shared environment for nearly 30 generations, and while the community composition has not shifted much among the bank vole lines, it is plausible that the bacterial metabolism within the host may have changed to adapt to the fast host metabolism including differential gene expression and food intake. Individual bacterial species may differ in their metabolic flexibility and sensitivity to environmental changes. Some species are able to switch energy sources depending on their availability (Porter et al. 2018) or rapidly evolve due to dietary changes (Dapa et al. 2022). The effect of dietary restriction on bacterial taxa is debated, with some studies suggesting that functional changes are more important than compositional ones, as different communities can produce the same functions (Kern et al. 2023). However, these effects remain poorly studied, and the influence of CR on microbiota resilience and plasticity is largely unknown.

Changes in differential abundance of microbiota taxa is a common response to dietary restriction in humans and rodent models (Kern et al. 2023). In previous studies, taxa such as *Bacteroides*, *Clostridium*, and *Lactobacillus* increased their proportions in response to dietary restriction, whereas *Helicobacter*, *Ruminococcaceae*, and *Coprococcus* decrease in relative abundance, and mixed effects are reported for *Bifidobacterium* and *Lachnospiraceae* (Kern et al. 2023). For example, switching to a diet rich in fiber / resistant starch results in significant shifts in microbial composition within 24 to 48 hours, primarily through the growth of fiber-degrading bacteria such as *Bifidobacterium*,

*Lactobacillus*, or *Ruminococcus* (Oliver et al. 2021; Wimmer et al. 2025). In our study, the *Bifidobacteriaceae* family was relatively more abundant in the SD diet, perhaps reflecting the more heterogeneous fiber composition of the SD diet compared to the CR diet, as these bacteria can utilize diverse carbohydrates (Lugli et al. 2017). In contrast, an earlier study using the same bank vole model found that *Lactobacillus* and *Bifidobacterium* increased in lines evolved to cope with a low-quality diet, whereas short-term exposure to the same diet for 10 days prior to sampling reduced their abundance (Lipowska et al. 2025). Together, these findings suggest that a combination of diet composition, host adaptation, and exposure history shape the observed responses of gut bacteria.

While cellulose supplementation reduces diet energy content, it can induce beneficial functional changes, for example by improving gut motility or SCFA production (Xie et al. 2022; Uddin et al. 2023) (but see also (Wen et al. 2022)). Our method of restricting energy intake with cellulose dilution may partially explain the discrepancy of our results compared to other studies applying caloric or dietary restriction (Kern et al. 2023). For example, in apparent contrast to previous studies, we observed a significant reduction in *Christensenellaceae*, one of the most abundant families in our data, in the CR treatment. Unlike nearly all other affected taxa, the reduction in *Christensenellaceae* lingered for at least two weeks after the diet restriction ended. Intriguingly, in human studies, this group of bacteria has been consistently found to have an inverse relationship with host body mass index and a positive association with metabolic health (Waters and Ley 2019). The finding that bacteria implicated in obesity and metabolic diseases are also affected by dietary restriction may help identify pathways of interest for therapeutic interventions.

Another notable finding from our study was that the *Helicobacter* genus and *Helicobacteraceae* family responded differently to dietary restriction depending on host metabolic background. Specifically, caloric restriction increased *Helicobacteraceae* in control animals but not in fast-metabolism lines, where abundances were already elevated under standard diet and did not further increase under diet restriction. *Helicobacteraceae* is typically present at low proportions in healthy rodent gut microbiota (e.g. 2% (A. Li et al. 2021)), but increase in abundance when the microbiota is disrupted through dietary (e.g. high fat diet) or metabolic challenges or diseases (X. Li et al. 2025; Péré-Védrenne et al. 2017; A. Li et al. 2021; Z. Zhao et al. 2022). Consequently, it is frequently considered a pathogenic group, including an important human pathogen *Helicobacter pylori* (Péré-Védrenne et al. 2017). Yet, one study suggests protective effects of non-pathogenic *Helicobacteraceae* against opportunistic pathogens in wild mice (B. Zhao et al. 2023). The role of the group of bacteria in our model system is not known, but the response may suggest opportunistic enrichment of the group under metabolic stress.

The digestive tract exhibits considerable plasticity and enlarges in response to increased food or fiber intake (Green and Millar 1987; Woodal 1989; Naya et al. 2007). In rodents, indigestible fiber such as cellulose can accumulate in the caecum, increasing digestive tract size and potentially affecting the microbial environment through changes in gut morphology and passage time (Overstreet et al. 2012; Wostmann 1981). Because fiber also binds water, it may increase apparent body mass and digestive organ size, potentially masking some effects of dietary restriction on body mass (Högberg and Lindberg 2004). Although our longitudinal design did not allow us to quantify transient morphological adjustments in the bank vole gut, the absence of differences in digestive tract mass between diet treatments at the end of the recovery period (P3) suggests that these changes were reversible. Such changes in metabolically active tissues may also contribute to the reduction in resting metabolic rate commonly observed under dietary restriction, including in our study (Norin and Metcalfe 2019) (Munch 1995; Kristan and Hammond 2001), but see also (R. M. Anderson and Weindruch 2007)). A-line voles nevertheless tended to have heavier digestive tracts, consistent with their higher body mass and food intake.

Several limitations should be acknowledged. First, our conclusions emphasize compositional shifts in microbiota, although functional outcomes such as energy harvest may depend more on changes in total bacterial abundance than on taxonomic composition, highlighting the need for quantitative and functional approaches (e.g., absolute bacterial loads, and/or metabolomics). Reliable functional assessment in non-model species is challenging (and was omitted here) because functional predictions depend on taxonomic assignment and genome annotations, which are largely lacking for poorly characterized microbial taxa. Second, the adaptive significance of these shifts was not directly tested; demonstrating this would require e.g. microbiota transplants into gnotobiotic hosts to assess effects on energy harvest efficiency. Finally, the study was conducted under relatively benign laboratory conditions, which may underestimate differences among lines; previous work (e.g. (Hanhimäki et al. 2022)) suggests that divergence in microbiota becomes more pronounced in semi-natural environments, indicating that future studies should incorporate ecologically more relevant stressors.

## Conclusion

Our results contribute experimental evidence on the interplay of host metabolic characteristics and gut microbiota properties in responding to dietary changes. While the gut microbiota can help buffer fluctuations in food availability, host traits may contribute to the type of response. The microbiota of high-metabolism voles showed a somewhat subdued response compared to control lines, suggesting stronger host regulation in animals subjected to prolonged directional selection. Stronger host control of the microbiota could stabilize host-microbiota interactions under stable conditions, but might constrain adaptability in fluctuating environments by limiting microbiota plasticity. An exploration of the responses of specific bacterial taxa to dietary challenges and the community consequences are a promising field for follow-up research.

## Author contributions

Designed the study: AMH, ES, PK, PW

Collected data: AMH, ES, K, ML, TI

Designed analyses: AMH, JKB, ML, PK, PL

Analysed data: AMH, JKB, PK

Wrote the paper: AMH, JKB

Approved the final version of the paper: all authors

## Data accessibility

Raw data will be made openly available upon acceptance of the manuscript for peer-reviewed publication.

## Acknowledgements

Special thanks to the team of technicians, students and assistants making this work possible in the midst of the first wave of COVID19, especially Anna Majtyka, Barbara Bober-Sowa, Natalia Strzelczyk, Katarzyna Baliga-Klimczyk, Karolina Sorys, Roksana Walkiewicz, Jerzy Skrobek, and Patrycja Maciak. Thanks to Alaa Hseiky for help with processing the RMR data, Monika Prus-Frankowska and Mateusz Buczek for advise on sample processing, and Watts/Mappes group for hosting in Jyväskylä.

## References

Amato, Katherine R., Steven R. Leigh, Angela Kent, et al. 2015. ‘The Gut Microbiota Appears to Compensate for Seasonal Diet Variation in the Wild Black Howler Monkey (Alouatta Pigra)’. Microbial Ecology 69 (2): 434–43. 10.1007/s00248-014-0554-7.

Anderson, Erik M., Jared M. Rozowsky, Brian J. Fazzone, et al. 2022. ‘Temporal Dynamics of the Intestinal Microbiome Following Short-Term Dietary Restriction’. Nutrients 14 (14): 14. 10.3390/nu14142785.

Anderson, Rozalyn M., and Richard Weindruch. 2007. ‘Metabolic Reprogramming in Dietary Restriction’. Interdisciplinary Topics in Gerontology 35: 18–38. 10.1159/000096554.

Baniel, Alice, Katherine R. Amato, Jacinta C. Beehner, et al. 2021. ‘Seasonal Shifts in the Gut Microbiome Indicate Plastic Responses to Diet in Wild Geladas’. Microbiome 9 (1). 10.1186/s40168-020-00977-9.

Bates, Douglas, Martin Mächler, Ben Bolker, and Steve Walker. 2015. ‘Fitting Linear Mixed-Effects Models Using Lme4’. Journal of Statistical Software 67 (October): 1–48. 10.18637/jss.v067.i01.

Besten, Gijs den, Karen van Eunen, Albert K. Groen, Koen Venema, Dirk-Jan Reijngoud, and Barbara M. Bakker. 2013. ‘The Role of Short-Chain Fatty Acids in the Interplay between Diet, Gut Microbiota, and Host Energy Metabolism’. Journal of Lipid Research 54 (9): 2325–40. 10.1194/jlr.R036012.

Bokulich, Nicholas A., Matthew R. Dillon, Yilong Zhang, et al. 2018. ‘Q2-Longitudinal: Longitudinal and Paired-Sample Analyses of Microbiome Data’. mSystems 3 (6): 10.1128/msystems.00219-18. 10.1128/msystems.00219-18.

Bolnick, Daniel I., Lisa K. Snowberg, J. Gregory Caporaso, Chris Lauber, Rob Knight, and William E. Stutz. 2014. ‘Major Histocompatibility Complex Class IIb Polymorphism Influences Gut Microbiota Composition and Diversity’. Molecular Ecology 23 (19): 4831–45. 10.1111/mec.12846.

Bolyen, Evan, Jai Ram Rideout, Matthew R. Dillon, et al. 2019. ‘Reproducible, Interactive, Scalable and Extensible Microbiome Data Science Using QIIME 2’. Nature Biotechnology 37 (8): 852–57. 10.1038/s41587-019-0209-9.

Bourdeau-Julien, Isabelle, Sophie Castonguay-Paradis, Gabrielle Rochefort, et al. 2023. ‘The Diet Rapidly and Differentially Affects the Gut Microbiota and Host Lipid Mediators in a Healthy Population’. Microbiome 11 (1): 26. 10.1186/s40168-023-01469-2.

Brandl, Simon J., Jonathan S. Lefcheck, Amanda E. Bates, Douglas B. Rasher, and Tommy Norin. 2023. ‘Can Metabolic Traits Explain Animal Community Assembly and Functioning?’ Biological Reviews 98 (1): 1–18. 10.1111/brv.12892.

Brooks, Mollie, E., Kasper Kristensen, Koen Benthem J.,van, et al. 2017. ‘glmmTMB Balances Speed and Flexibility Among Packages for Zero-Inflated Generalized Linear Mixed Modeling’. The R Journal 9 (2): 378. 10.32614/RJ-2017-066.

Brzęk, Pawel, Aneta Książek, Agnieszka Dobrzyn, and Marek Konarzewski. 2012. ‘Effect of Dietary Restriction on Metabolic, Anatomic and Molecular Traits in Mice Depends on the Initial Level of Basal Metabolic Rate’. Journal of Experimental Biology 215 (18): 3191–99. 10.1242/jeb.065318.

Carmody, Rachel N., Georg K. Gerber, Jesus M. Luevano, et al. 2015. ‘Diet Dominates Host Genotype in Shaping the Murine Gut Microbiota’. Cell Host & Microbe 17 (1): 72–84. 10.1016/j.chom.2014.11.010.

Chaston, John M., Adam J. Dobson, Peter D. Newell, and Angela E. Douglas. 2016. ‘Host Genetic Control of the Microbiota Mediates the Drosophila Nutritional Phenotype’. Applied and Environmental Microbiology 82 (2): 671–79. 10.1128/AEM.03301-15.

Dalby, Matthew J. 2023. ‘Questioning the Foundations of the Gut Microbiota and Obesity’. Philosophical Transactions of the Royal Society B: Biological Sciences 378 (1888): 20220221. 10.1098/rstb.2022.0221.

Dapa, Tanja, Ricardo Serotte Ramiro, Miguel Filipe Pedro, Isabel Gordo, and Karina Bivar Xavier. 2022. ‘Diet Leaves a Genetic Signature in a Keystone Member of the Gut Microbiota’. Cell Host & Microbe 30 (2): 183–199.e10. 10.1016/j.chom.2022.01.002.

David, Lawrence A., Corinne F. Maurice, Rachel N. Carmody, et al. 2014. ‘Diet Rapidly and Reproducibly Alters the Human Gut Microbiome’. Nature 505 (7484): 559–63. 10.1038/nature12820.

Davis, Nicole M., Diana M. Proctor, Susan P. Holmes, David A. Relman, J. Benjamin, and William Moore Drive. 2018. Simple Statistical Identification and Removal of Contaminant Sequences in Marker-Gene and Metagenomics Data. no. 919: 1–39.

Dheyongera, Geoffrey, Katherine Grzebyk, Agata M. Rudolf, Edyta T. Sadowska, and Paweł Koteja. 2016. ‘The Effect of Chlorpyrifos on Thermogenic Capacity of Bank Voles Selected for Increased Aerobic Exercise Metabolism’. Chemosphere 149 (April): 383–90. 10.1016/j.chemosphere.2015.12.120.

Faulks, Sally C., Nigel Turner, Paul L. Else, and A. J. Hulbert. 2006. ‘Calorie Restriction in Mice: Effects on Body Composition, Daily Activity, Metabolic Rate, Mitochondrial Reactive Oxygen Species Production, and Membrane Fatty Acid Composition’. The Journals of Gerontology: Series A 61 (8): 781–94. 10.1093/gerona/61.8.781.

Glazier, Douglas S., and Vojsava Gjoni. 2024. ‘Interactive Effects of Intrinsic and Extrinsic Factors on Metabolic Rate’. Philosophical Transactions of the Royal Society B: Biological Sciences 379 (1896): 20220489. 10.1098/rstb.2022.0489.

Green, D. A., and J. S. Millar. 1987. ‘Changes in Gut Dimension and Capacity of Peromyscus Maniculatus Relative to Diet Quality and Energy Needs’. Can. J. Zool. 65: 2159–62.

Grosiak, Marta, Paweł Koteja, Ulf Bauchinger, and Edyta T. Sadowska. 2020. ‘Age-Related Changes in the Thermoregulatory Properties in Bank Voles From a Selection Experiment’. Frontiers in Physiology 11 (November). 10.3389/fphys.2020.576304.

Hall, Charles A. S., Jack A. Stanford, and F. Richard Hauer. 1992. ‘The Distribution and Abundance of Organisms as a Consequence of Energy Balances along Multiple Environmental Gradients’. Oikos 65 (3): 377–90. 10.2307/3545553.

Hanhimäki, Elina, Phillip C. Watts, Esa Koskela, Paweł Koteja, Tapio Mappes, and Anni M. Hämäläinen. 2022. ‘Evolved High Aerobic Capacity Has Context-Specific Effects on Gut Microbiota’. Frontiers in Ecology and Evolution 10. https://www.frontiersin.org/articles/10.3389/fevo.2022.934164.

Heiss, Christina N., and Louise E. Olofsson. 2017. ‘Gut Microbiota-Dependent Modulation of Energy Metabolism’. Journal of Innate Immunity 10 (3): 163–71. 10.1159/000481519.

Högberg, Ann, and Jan Erik Lindberg. 2004. ‘Influence of Cereal Non-Starch Polysaccharides and Enzyme Supplementation on Digestion Site and Gut Environment in Weaned Piglets’. Animal Feed Science and Technology 116 (1): 113–28. 10.1016/j.anifeedsci.2004.03.010.

Hseiky, Alaa, Małgorzata M. Lipowska, Edyta T. Sadowska, Alicja Józkowicz, Witold N. Nowak, and Paweł Koteja. 2025. ‘Effect of Western Diet on Body Composition, Locomotor Performance and Blood Biochemical Profile in the Bank Vole’. Journal of Experimental Biology 228 (19): jeb250698. 10.1242/jeb.250698.

Hseiky, Alaa, Edyta T. Sadowska, and Paweł Koteja. 2026. ‘The Effects of Short-Term Consumption of a Western Diet on Aerobic Exercise Performance in Bank Voles with Inherently Distinct Metabolic Rates’. Experimental Physiology 111 (3): 930–45. 10.1113/EP092646.

Huang, Ce, Shengyu Feng, Fengjiao Huo, and Hailiang Liu. 2022. ‘Effects of Four Antibiotics on the Diversity of the Intestinal Microbiota’. Microbiology Spectrum 10 (2): e01904–21. 10.1128/spectrum.01904-21.

Jaromin, Ewa, Edyta T. Sadowska, and Paweł Koteja. 2019. ‘Is Experimental Evolution of an Increased Aerobic Exercise Performance in Bank Voles Mediated by Endocannabinoid Signaling Pathway?’ Frontiers in Physiology 10 (May). 10.3389/fphys.2019.00640.

Kern, Lara, Denise Kviatcovsky, Yiming He, and Eran Elinav. 2023. ‘Impact of Caloric Restriction on the Gut Microbiota’. Current Opinion in Microbiology 73 (June): 102287. 10.1016/j.mib.2023.102287.

Klimenko, Natalia S., Vera E. Odintsova, Anastasia Revel-Muroz, and Alexander V. Tyakht. 2022. ‘The Hallmarks of Dietary Intervention-Resilient Gut Microbiome’. Npj Biofilms and Microbiomes 8 (1): 1–11. 10.1038/s41522-022-00342-8.

Kohl, Kevin D., Edyta T. Sadowska, Agata M. Rudolf, M. Denise Dearing, and Paweł Koteja. 2016. ‘Experimental Evolution on a Wild Mammal Species Results in Modifications of Gut Microbial Communities’. Frontiers in Microbiology 7. 10.3389/fmicb.2016.00634.

Kolodny, Oren, and Hinrich Schulenburg. 2020. ‘Microbiome-Mediated Plasticity Directs Host Evolution along Several Distinct Time Scales’. Philosophical Transactions of the Royal Society B: Biological Sciences 375 (1808): 20190589. 10.1098/rstb.2019.0589.

Korach-Rechtman, Hila, Shay Freilich, Shiran Gerassy-Vainberg, et al. 2019. ‘Murine Genetic Background Has a Stronger Impact on the Composition of the Gut Microbiota than Maternal Inoculation or Exposure to Unlike Exogenous Microbiota’. Applied and Environmental Microbiology 85 (18): e00826–19. 10.1128/AEM.00826-19.

Kristan, Deborah M., and Kimberly A. Hammond. 2001. ‘Parasite Infection and Caloric Restriction Induce Physiological and Morphological Plasticity’. *American Journal of Physiology-Regulatory*, Integrative and Comparative Physiology 281 (2): R502–10. 10.1152/ajpregu.2001.281.2.R502.

Lenth, Russell V. 2025. ‘Emmeans: Estimated Marginal Means, Aka Least-Squares Means’. *R Package Version 1.10.7*. https://CRAN.R-project.org/package=emmeans.

Li, Aili, Nana Wang, Na Li, et al. 2021. ‘Modulation Effect of Chenpi Extract on Gut Microbiota in High-Fat Diet-Induced Obese C57BL/6 Mice’. Journal of Food Biochemistry 45 (4): e13541. 10.1111/jfbc.13541.

Li, Xin, Yanting Qin, Fangfang Yue, and Xin Lü. 2025. ‘Comprehensive Analysis of Fecal Microbiome and Metabolomics Uncovered Dl-Norvaline-Ameliorated Obesity-Associated Disorders in High-Fat Diet-Fed Obese Mice by Targeting the Gut Microbiota’. Journal of Agricultural and Food Chemistry 73 (4): 2381–92. 10.1021/acs.jafc.4c06638.

Lin, Huang, Merete Eggesbø, and Shyamal Das Peddada. 2022. ‘Linear and Nonlinear Correlation Estimators Unveil Undescribed Taxa Interactions in Microbiome Data’. Nature Communications 13 (1): 4946. 10.1038/s41467-022-32243-x.

Lin, Huang, and Shyamal Das Peddada. 2020. ‘Analysis of Compositions of Microbiomes with Bias Correction’. Nature Communications 11 (1): 3514. 10.1038/s41467-020-17041-7.

Lin, Huang, and Shyamal Das Peddada. 2024. ‘Multigroup Analysis of Compositions of Microbiomes with Covariate Adjustments and Repeated Measures’. Nature Methods 21 (1): 83–91. 10.1038/s41592-023-02092-7.

Lindsay, Elle C., Neil B. Metcalfe, and Martin S. Llewellyn. 2020. ‘The Potential Role of the Gut Microbiota in Shaping Host Energetics and Metabolic Rate’. Journal of Animal Ecology 89 (11): 2415–26. 10.1111/1365-2656.13327.

Lipowska, Małgorzata M., Geoffrey Dheyongera, Edyta T. Sadowska, and Paweł Koteja. 2019. ‘Experimental Evolution of Aerobic Exercise Performance and Hematological Traits in Bank Voles’. Comparative Biochemistry and Physiology Part A: Molecular & Integrative Physiology 234 (August): 1–9. 10.1016/j.cbpa.2019.04.008.

Lipowska, Małgorzata M., Edyta T. Sadowska, Ulf Bauchinger, Wolfgang Goymann, Barbara Bober-Sowa, and Paweł Koteja. 2020. ‘Does Selection for Behavioral and Physiological Performance Traits Alter Glucocorticoid Responsiveness in Bank Voles?’ Journal of Experimental Biology 223 (15): jeb219865. 10.1242/jeb.219865.

Lipowska, Małgorzata M., Edyta T. Sadowska, Kevin D. Kohl, and Paweł Koteja. 2025. ‘Experimental Evolution of a Mammalian Holobiont: Bank Voles Selected for Herbivorous Capability Evolved Distinct and Robust Gut Bacterial Communities’. ISME Communications 5 (1): ycaf160. 10.1093/ismeco/ycaf160.

Low, Adrian, Melissa Soh, Sou Miyake, et al. 2021. ‘Longitudinal Changes in Diet Cause Repeatable and Largely Reversible Shifts in Gut Microbial Communities of Laboratory Mice and Are Observed across Segments of the Entire Intestinal Tract’. International Journal of Molecular Sciences 22 (11): 11. 10.3390/ijms22115981.

Lugli, Gabriele Andrea, Christian Milani, Francesca Turroni, et al. 2017. ‘Comparative Genomic and Phylogenomic Analyses of the Bifidobacteriaceae Family’. BMC Genomics 18 (1): 568. 10.1186/s12864-017-3955-4.

Lynch, J. B., and E. Y. Hsiao. 2019. ‘Microbiomes as Sources of Emergent Host Phenotypes’. Science 365 (6460): 1405–9. 10.1126/science.aay0240.

Maiti, Uttaran, Edyta T. Sadowska, Katarzyna M. ChrzĄścik, and Paweł Koteja. 2019. ‘Experimental Evolution of Personality Traits: Open-Field Exploration in Bank Voles from a Multidirectional Selection Experiment’. Current Zoology 65 (4): 375–84. 10.1093/cz/zoy068.

Martin, Corby K., Leonie K. Heilbronn, Lilian de Jonge, et al. 2007. ‘Effect of Calorie Restriction on Resting Metabolic Rate and Spontaneous Physical Activity’. Obesity 15 (12): 2964–73. 10.1038/oby.2007.354.

Munch, I. C. 1995. ‘Influences of Time Intervals between Meals and Total Food Intake on Resting Metabolic Rate in Rats’. Acta Physiologica Scandinavica 153 (3): 243–47. 10.1111/j.1748-1716.1995.tb09859.x.

Naya, Daniel E., William H. Karasov, and Francisco Bozinovic. 2007. ‘Phenotypic Plasticity in Laboratory Mice and Rats: A Meta-Analysis of Current Ideas on Gut Size Flexibility’. Evolutionary Ecology Research 9: 1363–74.

Norin, Tommy, and Neil B. Metcalfe. 2019. ‘Ecological and Evolutionary Consequences of Metabolic Rate Plasticity in Response to Environmental Change’. Philosophical Transactions of the Royal Society B: Biological Sciences 374 (1768): 20180180. 10.1098/rstb.2018.0180.

Nowak, Karol H., Emily Hartop, Monika Prus-Frankowska, et al. 2025. ‘What Lurks in the Dark? An Innovative Framework for Studying Diverse Wild Insect Microbiota’. Microbiome 13 (1): 186. 10.1186/s40168-025-02169-9.

Oliver, Andrew, Alexander B. Chase, Claudia Weihe, et al. 2021. ‘High-Fiber, Whole-Food Dietary Intervention Alters the Human Gut Microbiome but Not Fecal Short-Chain Fatty Acids’. mSystems 6 (2): 10.1128/msystems.00115-21. 10.1128/msystems.00115-21.

Overstreet, Anne-Marie C., Amanda E. Ramer-Tait, Albert E. Jergens, et al. 2012. ‘The Role of the Microbiota in Gastrointestinal Health and Disease’. In Inflammatory Bowel Disease. IntechOpen. 10.5772/53920.

Péré-Védrenne, Christelle, Bram Flahou, Mun Fai Loke, Armelle Ménard, and Jamuna Vadivelu. 2017. ‘Other Helicobacters, Gastric and Gut Microbiota’. Helicobacter 22 (S1): e12407. 10.1111/hel.12407.

Pinheiro, José, and Douglas Bates. 2000. Mixed-Effects Models in S and S-PLUS. Springer. 10.1007/b98882.

Pinheiro, José, Douglas Bates, and R Core Team. 2024. ‘Nlme: Linear and Nonlinear Mixed Effects Models’. *R Package Version 3.1-166*. https://CRAN.R-project.org/package=nlme.

Porter, Nathan T., Ana S. Luis, and Eric C. Martens. 2018. ‘Bacteroides Thetaiotaomicron’. Trends in Microbiology 26 (11): 966–67. 10.1016/j.tim.2018.08.005.

R Core Team. 2018. ‘R: A Language and Environment for Statistical Computing.’ Preprint, R Foundation for Statistical Computing.

Ramsey, Jon J., Mary-Ellen Harper, and Richard Weindruch. 2000. ‘Restriction of Energy Intake, Energy Expenditure, and Aging’. Free Radical Biology and Medicine 29 (10): 946–68. 10.1016/S0891-5849(00)00417-2.

Sadowska, Edyta T., Katarzyna Baliga-Klimczyk, Katarzyna M. Chrząścik, and Paweł Koteja. 2008. ‘Laboratory Model of Adaptive Radiation: A Selection Experiment in the Bank Vole’. Physiological and Biochemical Zoology 81 (5): 627–40. 10.1086/590164.

Sadowska, Edyta T., Clare Stawski, Agata Rudolf, et al. 2015a. ‘Evolution of Basal Metabolic Rate in Bank Voles from a Multidirectional Selection Experiment.’ Proceedings. Biological Sciences (England) 282 (1806): 20150025. 10.1098/rspb.2015.0025.

Sadowska, Edyta T., Clare Stawski, Agata Rudolf, et al. 2015b. ‘Evolution of Basal Metabolic Rate in Bank Voles from a Multidirectional Selection Experiment’. Proceedings of the Royal Society B: Biological Sciences 282 (1806): 20150025. 10.1098/rspb.2015.0025.

Salzman, Nita H., Kuiechun Hung, Dipica Haribhai, et al. 2010. ‘Enteric Defensins Are Essential Regulators of Intestinal Microbial Ecology’. Nature Immunology 11 (1): 76–82. 10.1038/ni.1825.

Selman, Colin, Tracey Phillips, Jessica L. Staib, Jackie S. Duncan, Christiaan Leeuwenburgh, and John R. Speakman. 2005. ‘Energy Expenditure of Calorically Restricted Rats Is Higher than Predicted from Their Altered Body Composition’. Mechanisms of Ageing and Development 126 (6): 783–93. 10.1016/j.mad.2005.02.004.

Smith, Reuben L., Maarten R. Soeters, Rob C. I. Wüst, and Riekelt H. Houtkooper. 2018. ‘Metabolic Flexibility as an Adaptation to Energy Resources and Requirements in Health and Disease’. Endocrine Reviews 39 (4): 489–517. 10.1210/er.2017-00211.

Sohal, Rajindar S., Melissa Ferguson, Barbara H. Sohal, and Michael J. Forster. 2009. ‘Life Span Extension in Mice by Food Restriction Depends on an Energy Imbalance12’. The Journal of Nutrition 139 (3): 533–39. 10.3945/jn.108.100313.

Spor, Aymé, Omry Koren, and Ruth Ley. 2011. ‘Unravelling the Effects of the Environment and Host Genotype on the Gut Microbiome’. Nature Reviews Microbiology 9 (4): 279–90. 10.1038/nrmicro2540.

Stamatakis, Alexandros, Jiajie Zhang, and Kassian Kobert. 2014. Genome Analysis PEAR : A Fast and Accurate Illumina Paired-End reAd mergeR. 30 (5): 614–20. 10.1093/bioinformatics/btt593.

Tremaroli, Valentina, and Fredrik Bäckhed. 2012. ‘Functional Interactions between the Gut Microbiota and Host Metabolism’. Nature 489 (7415): 242–49. 10.1038/nature11552.

Turnbaugh, Peter J., Ruth E. Ley, Michael A. Mahowald, Vincent Magrini, Elaine R. Mardis, and Jeffrey I. Gordon. 2006. ‘An Obesity-Associated Gut Microbiome with Increased Capacity for Energy Harvest’. Nature 444 (7122): 1027–31. 10.1038/nature05414.

Turnbaugh, Peter J., Vanessa K. Ridaura, Jeremiah J. Faith, Federico E. Rey, Rob Knight, and Jeffrey I. Gordon. 2009. ‘The Effect of Diet on the Human Gut Microbiome: A Metagenomic Analysis in Humanized Gnotobiotic Mice’. Science Translational Medicine 1 (6): 6ra14. 10.1126/scitranslmed.3000322.

Uddin, Md Karim, Md Rayhan Mahmud, Shah Hasan, Olli Peltoniemi, and Claudio Oliviero. 2023. ‘Dietary Micro-Fibrillated Cellulose Improves Growth, Reduces Diarrhea, Modulates Gut Microbiota, and Increases Butyrate Production in Post-Weaning Piglets’. Scientific Reports 13 (1): 6194. 10.1038/s41598-023-33291-z.

Voolstra, Christian R., and Maren Ziegler. 2020. ‘Adapting with Microbial Help: Microbiome Flexibility Facilitates Rapid Responses to Environmental Change’. BioEssays 42 (7): 2000004. 10.1002/bies.202000004.

Wang, Luanfeng, Fang Wang, Ling Xiong, Haizhao Song, Bo Ren, and Xinchun Shen. 2024. ‘A Nexus of Dietary Restriction and Gut Microbiota: Recent Insights into Metabolic Health’. Critical Reviews in Food Science and Nutrition 64 (24): 8649–71. 10.1080/10408398.2023.2202750.

Waters, Jillian L., and Ruth E. Ley. 2019. ‘The Human Gut Bacteria Christensenellaceae Are Widespread, Heritable, and Associated with Health’. BMC Biology 17 (1): 83. 10.1186/s12915-019-0699-4.

Wen, Song, Guifang Yuan, Cunya Li, Yang Xiong, Xuemei Zhong, and Xiaoyu Li. 2022. ‘High Cellulose Dietary Intake Relieves Asthma Inflammation through the Intestinal Microbiome in a Mouse Model’. PLOS ONE 17 (3): e0263762. 10.1371/journal.pone.0263762.

Wickham, Hadley. 2016. Ggplot2. Use R! Springer International Publishing. 10.1007/978-3-319-24277-4.

Wilde, Jacob, Emma Slack, and Kevin R. Foster. 2024. ‘Host Control of the Microbiome: Mechanisms, Evolution, and Disease’. Science 385 (6706). 10.1126/science.adi3338.

Wimmer, Benedikt H., Sarah Moraïs, Itai Amit, et al. 2025. ‘Spatial Constraints Drive Amylosome-Mediated Resistant Starch Degradation by Ruminococcus Bromii in the Human Colon’. Nature Communications 16 (1): 10763. 10.1038/s41467-025-65800-1.

Woodal, P. F. 1989. ‘The Effects of Increased Dietary Cellulose on the Anatomy, Physiology and Behaviour of the Captive Water Voles, Arvicola Terrestris (L.) (Rodentia: Microtinae)’. Comp. Biochem. Physiol. 94A: 615–21.

Wostmann, Bernard S. 1981. ‘THE GERMFREE ANIMAL IN NUTRITIONAL STUDIES’. Annual Review of Nutrition 1 (Volume 1, 1981): 257–79. 10.1146/annurev.nu.01.070181.001353.

Xie, Tao, Fa Jin, Xiaokun Jia, Hengxu Mao, Yuting Xu, and Shizhong Zhang. 2022. ‘High Cellulose Diet Promotes Intestinal Motility through Regulating Intestinal Immune Homeostasis and Serotonin Biosynthesis’. Biological Chemistry 403 (3): 279–92. 10.1515/hsz-2021-0216.

Zhao, Bei, Lisa Osbelt, Till Robin Lesker, et al. 2023. ‘*Helicobacter* Spp. Are Prevalent in Wild Mice and Protect from Lethal *Citrobacter Rodentium* Infection in the Absence of Adaptive Immunity’. Cell Reports 42 (6): 112549. 10.1016/j.celrep.2023.112549.

Zhao, Zhifang, Dejun Cui, Guosong Wu, et al. 2022. ‘Disrupted Gut Microbiota Aggravates Working Memory Dysfunction Induced by High-Altitude Exposure in Mice’. Frontiers in Microbiology 13 (November). 10.3389/fmicb.2022.1054504.

