## Supplementary information for "Selection for high metabolic rate reduces gut microbiota responsiveness to dietary restriction in bank voles"

### Supplementary materials and methods

#### Study design

In total, 224 adult voles from the 29th generation of the selection experiment were selected for the experiment at age 4-5 months and randomly allocated to two alternative diet treatment groups, such that 7 voles / sex / replicate line / selection line type / diet treatment were enrolled in the experiment. Due to time and space limitations the experiment was performed in 14 overlapping blocks (started on consecutive days), with 14-21 individuals per block. The animals were distributed randomly with respect to the cage position in the ventilation system.

Upon enrollment, the voles were housed singly in individually ventilated cages (IVC) (Techniplast, GM500, Italy) to minimize transmission of gut bacteria among individuals during the experiment. Each cage was provided with a shelter (cleaned coconut shell) and dust-free aspen bedding (Abedd, Germany). The voles were maintained under ambient temperature (21° C) and photoperiod (16:8 h light/dark), with water and food (according to each individual's assigned diet regime) available *ad libitum* throughout the experiment. The experiment began with 224 individuals, of which 12 individuals died during the study (not biased for diet treatment or selection line type).

#### Sampling and measurement techniques

##### Food consumption

Individual food consumption and calorie intake were approximated throughout the experiment. For this, 30-50 g of the appropriate diet (SD, CR15 or CR30 according to the experimental phase and treatment group) was weighed to ±0.01 g and placed into the feeder of each animal. After 3-4 days the remaining food was removed, oven-dried (60C), and weighed (±0.001 g, Radwag PS 200/2000.R2 Precision Balance, Radom, Poland). To ensure animals had ad lib access to food, the presence of food in the feeder was checked daily and supplemented with a known amount as necessary. Average daily food consumption was approximated as the difference in the mass of food provided and recovered divided by the number of days. This approach was chosen over more accurate ways of measuring food consumption (e.g. using a grid flooring without bedding in the cages) for reasons of animal welfare and prioritizing repeated measurements. Although we took care to search the bedding for (rare) pellet fragments, it is likely that some fragments were missed, thus the food consumption should not be interpreted in absolute but relative quantities. We have no reason to think that the method causes any major bias towards either selection direction or diet type.

#### Resting metabolic rate

Resting metabolic rate (RMR) was measured similarly to our previous measurements of basal metabolic rate (Sadowska et al., 2015)(Hseiky et al. 205), except that the measurements were performed at the regular housing temperature of 20°C. Briefly, the animals were weighed and placed in plastic respirometric chambers 850 ml) without food or water. The chambers (850 ml) were placed in a dark, climate-controlled room at 20°C. The rate of oxygen consumption ($VO_2$; ml $O_2$ min$^{-1}$) was measured using an eight-channel, open-flow, positive-pressure respirometric system.

Fresh air from outside the room was dried with silica gel and pumped into the eight chambers: seven containing animals and one empty reference chamber. The flow rate was stabilized at a nominal 450 ml min$^{-1}$ with GFC-17 thermal mass-flow controllers (AALBORG, Orangeburg, NY, USA), separately for each channel. The actual flow rate in each channel (407 – 451 ml min$^{-1}$ STPD) was calculated after calibrating the mass-flow controllers against a precise LO 63/33 rotameter (Rota, Germany). Samples of air flowing out of the eight chambers (approximately 150 ml min$^{-1}$) were taken sequentially in a 13-min cycle using an RM-8 multiplexer (Sable Systems, Las Vegas, NV, USA). The air samples were dried with an ND2 Nafion-tube water vapor absorber (Sable Systems, Las Vegas, NV, USA), followed by magnesium perchloride, and directed to a $CO_2$ analyzer (LI-850, LI-COR Biosciences, Lincoln, NE, USA) and an oxygen analyzer (AMETEK S3AII, Applied Electrochemistry, Pittsburgh, PA, USA). Mean values of the analogue outputs from the $O_2$ and $CO_2$ analyzers were recorded every 4 s using a UE-9 AD interface (LabJack Corporation, Lakewood, CO, USA) and a custom-made protocol implemented in the DAQFactory acquisition system (Azeotech, Ashland, OR, USA). $VO_2$ was calculated from values recorded during the last 20 s before switching to the next channel, according to the appropriate respirometric equation (Sadowska et al., 2015). Because the half-time of air exchange in the chambers exceeded 2 min (850/420 ≈ 2.0 min), the 20-s measurements reflected signals integrated over several minutes. Movement activity of the animals and the background vibration recorded in the empty reference chamber were monitored continuously with MAD-1 gravimetric detectors (Sable Systems, Las Vegas, NV, USA).

The measurements were planned to last 4.5 h (plus up to 30 min for weighing the animals and preparing the system), which would allow 20 measurement cycles to be recorded, each including the reference chamber and the seven chambers containing animals. However, because of logistical constraints (the need to perform procedures simultaneously on other groups of animals) and technical problems (such as animals escaping during weighing and instability of the respirometric system), the interval between weighing the animals and the start of the recordings ranged from 5 to

130 min (mean ± SD: 30 ± 22 min). For the same reasons, the recording time varied from 215 to 320 min (mean ± SD: 252 ± 13 min), allowing $VO_2$ to be calculated for 16 – 24 cycles. The first two cycles were always excluded from the analyses, and in some cases additional cycles at the beginning or end of a trial were excluded because of unstable baseline values. Because values obtained during the final part of very long trials would not be comparable with those from most other trials, the data were truncated to a maximum of 19 cycles (effectively 17 cycles, because the first two cycles were always excluded). RMR was operationally defined as the mean value of the three lowest $VO_2$ values recorded for a given individual.

#### Body composition and organ mass

On the last day of the experiment (Day 48), the voles were euthanized and dissected to quantify differences in body fat content and digestive tract mass among treatments. Voles were weighed to ±0.01g, euthanized with isoflurane inhalation overdose, and the digestive tract from stomach to anus was dissected. Any visible visceral fat was carefully removed and placed back with the carcass. Stomach, caecum and intestine (large + small intestine) were dissected separately, placed in saline solution, carefully emptied of contents, freeze dried (Beta 1-8 LD plus; Christ, Osterode, Germany) and weighed to ±0.001 g to obtain dry organ masses.

Total body fat was extracted from entire bodies (excluding digestive tract). The carcasses were freeze-dried (Beta 1-8 LD plus; Christ, Osterode, Germany) to a constant mass (±0.01g). The dry weight of the bodies was measured to ±0.001 g resolution, pulverized using a food grinder (COSO Design) and total body fat was extracted by hot Soxhlet extraction using a four-column BÜCHI extractor (Operation Manual Extraction System B-811 / B-811 LSV, BÜCHI Labortechnik AG, Flawil, Switzerland. The extracted product was dried and weighed and body fat percentage was calculated as the difference in mass before and after fat extraction.

For a subset of animals, dissections could not be performed on day 48 due to unforeseen COVID19-restrictions and were delayed by one week. These individuals were subjected again to the RMR measurement protocol (data not shown) the day before dissection to match the conditions prior to dissection. The delay in dissection is controlled for in the models of intestine mass and body fat content.

#### Fecal sampling

For fecal microbiota analysis, fecal samples (at P1-P3) were collected by placing an individual into a clean empty cage, the floor of which had been wiped with ethanol, for 90 minutes, with *ad libitum* access to water. At the end of this time, the individual was removed and fecal pellets collected from the cage floor using sterilized tools and avoiding urine contamination. Samples were placed immediately on ice and transferred to -30℃ freezer for max. 24 h, after which they were stored at -80 ℃.

#### DNA extraction and sequencing

We selected for sequencing samples from vole individuals for which we had the most complete data on variables of interest, balancing treatment groups, sexes, and selection lines. Total genomic DNA was extracted from 480 fecal samples from 169 individuals at three different time points, at the end of each three phases of the experiment: P1 (baseline), P2 (diet treatment), and P3 (recovery). Additionally, to account for possible contamination of samples by reagents or laboratory equipment, negative controls were collected of both isolation and library preparation phases (total 12 control samples). The fecal samples were thawed on ice before DNA extraction. Total genomic DNA was isolated from ca. 100 ug sample of feces using the same batch of DNeasy Power Soil Pro kit (Qiagen, Germany) for all samples in aseptic conditions using the manufacturer's instructions. The purity and concentration were measured using Nanodrop.

To obtain libraries of bacteria genetic material, we used a two-step PCR library preparation protocol to multiply the variants of the 16S rRNA gene V4 region in accordance with the Earth Microbiome Project guidelines (as in Glenn et al. 2019; Marquina et al. 2021). First, the V4 target region was amplified using custom 515F and 806R primers with variable-length inserts and Illumina adapter tails to ensure that every sample had a unique index. The products were purified using in-house produced SPRI (solid phase reversible immobilization) magnetic beads. Then, the samples were indexed using unique combinations of a custom set of 192 forward and 192 reverse indexing primers (Iwaszkiewicz-Eggebrecht et al. 2023).

V4 regions were amplified in PCR reaction, using 515F and 806R 16S rRNA gene primers containing 0-3b-long insert sequences at 5'-ends. Reaction was performed on 96-well plates, using 5ul Quiagen Multiplex MasterMix, 1ul 10mM solution of each of the primers, 2ul of ddH2O and 1ul of DNA dilution. The reaction was performed for 22 cycles, with a program commonly used by Piotr Łukasik's team to amplify rRNA gene regions (94C-30s, 50C-19s, 72C-90s). Effects of the reaction were checked by running 2ul of each sample on electrophoresis gel. If the resulting band was weaker than expected, PCR was repeated for this sample. PCR plates were stored in a fridge (short-term) and in -20C (long-term).

PCR products were cleaned with SPRI magnetic beads produced by the Piotr Łukasik's team, using the entire volume of PCR products (8ul) and 16ul of beads. The beads were washed twice with 80% EtOH and DNA was eluted with 20.5ul of TE buffer. Elution efficiency was checked with electrophoresis. Plates containing cleaned PCR products were stored in a fridge (short-term) and in -20C (long-term).

Cleaned PCR products were individually marked by adding indexing sequences at both ends, with a second PCR reaction. Primers were sampled from Piotr Łukasik's custom-designed stock of 192 primers F and 192 primers R. An additional Neg sample (ddH20) was added to each plate and processed alongside with other samples. Each sample was marked with an unique combination of one primer F and one primer R, each primer was used in no more than 8 combinations. The PCR mix consisted of 5ul Quiagen Multiplex MasterMix, 1ul 10mM solution of each of the primers, 2ul of ddH2O and 1ul of DNA, the reaction was run for 7 cycles using the same program as that for V4 region amplification. Effects of the reaction were checked by running 2ul of each sample on electrophoresis gel. If the resulting band was weaker than expected, PCR was repeated for this sample. PCR plates were stored in a fridge (short-term) and in -20C (long-term).

#### Read data processing

Demultiplexed forward and reverse reads had primers trimmed using a custom script https://github.com/Symbiosis-JU/Bioinformatic-pipelines/blob/main/multiPISS.py, and were then assembled into contigs using PEAR (Stamatakis et al. 2014), which merges reads by maximizing the assembly score of the read overlap. The cutoff threshold was set to 260 bp since it involved 99.9% of sequences. Further analyses were performed in QIIME2 version 2023.5.1 (Bolyen et al. 2019). Initially, the total number of forward reads equaled 44,853,914 and decreased by 1.1% to 44,375,771 (478,143 reads discarded) after an initial quality filtering process based on quality scores (quality threshold 30). Reads were denoised and chimera removed using deblur and trimmed to 252 bp, resulting in the rejection of nine samples (including eight extraction blanks and one vole sample) that included only reads shorter than this threshold. The resulting feature table consisted of 18,974,257 sequences with 4,012 amplicon sequence variants (ASVs). Taxonomy was assigned using a naïve Bayes classifier (QIIME2 sklearn) that has been pretrained on the Silva version 138 16S rRNA database. After adding taxonomy, the feature table was cleaned from archaea, chloroplasts and mitochondria.

To identify and remove possible contaminant sequences, we examined the relative frequency and prevalence of ASVs in contamination control samples (extraction blanks) relative to the fecal samples. For this, we imported the data into program R (version 4.2.2 (R Core Team 2018)) as a phyloseq (McMurdie and Holmes 2013) object and then cleaned using package decontam (Davis et al. 2018).

The contaminant features were identified using both prevalence and frequency methods with default parameters, however, for further steps only the sequences identified by frequency method were used. Altogether, 29 ASVs were considered as contaminants due to their higher frequency in the contamination controls than in fecal samples. These ASVs were excluded from the microbiome data, leaving 3,960 ASVs. The contamination control samples and contaminant sequences were discarded from the data set, and the data were re-imported into QIIME2. The remaining sequences were classified again to assign taxonomy. Data was rarefied to 10,058 sequences per sample and this normalized feature table was used for further analyses. During rarefaction 4 samples were dropped, leaving 474 samples (from 168 voles; N= 158 + 159 + 157 samples from phases P1-P3, respectively) with 4,767,492 (25.91%) features as the final data set for downstream analyses of microbiota.

Alpha diversity of the gut microbiota was estimated by calculating ASV richness and Shannon index for each sample using rarefied data with R package phyloseq (version 1.42.0 (McMurdie and Holmes 2013)). Pairwise sample dissimilarities were calculated using three different beta diversity metrics (Bray–Curtis dissimilarity, unweighted and weighted UniFrac distances). Sample clustering patterns were visualized by principal coordinates analysis (PCoA) (McMurdie and Holmes 2013).

To quantify the magnitude of longitudinal (within-individual) changes in the diversity and community composition, "first distances" were extracted from beta diversity distance matrices in QIIME2. The first distances quantify the magnitude of change from one time point to another, i.e. distance between the position of two samples in a multidimensional distance matrix. We calculated the magnitude of change from baseline P1 to P2 (effect of diet treatment on microbiota, i.e. resistance), and from P1 to P3 (completeness of recovery, i.e. resilience). These measurements were imported into R for statistical analyses (see below).

#### Statistical analyses

Random slopes were initially included for replicate selection line (C1-4, A1-4) against diet treatment, phase, and sex to account for possible differences in the magnitude of the responses among replicate selection lines, but as these complex structures frequently had zero variance, prevented model convergence, and destabilized the models, a random intercept of vole identity nested within replicate line was chosen for all models.

For host traits, we used a compound symmetry model to accommodate residual covariances owing to autocorrelation of repeated measures across experimental phases. We initially compared fully unstructured, autoregressive, and compound symmetry approaches. Models with fully unstructured residual covariance were not reliably estimable due to incomplete repeated measures and limited observations per individual. To account for repeated measurements within individuals, we evaluated compound symmetry and autoregressive (AR(1)) residual covariance structures. Because multiple observations could occur within a given experimental phase and not all phases were observed for each individual, AR(1) structures were specified over observation order rather than explicit phase values. Results were qualitatively consistent across covariance structures. Therefore, for parsimony, the compound symmetry model was retained for primary inference. Where estimable, AR(1) and compound symmetry residual structures yielded nearly identical model fits (ΔAIC < 2).

We also estimated a separate residual variance for each phase with weights = varIdent(~ 1 | Phase). in all models excepting the body mass model, where limited within-phase variability prevented the estimation, as the combination of structures was not identifiable for some groups. A homogeneous residual variance structure was thus retained the body mass model.

The absence of significant multicollinearity was confirmed by computing predictor variance inflation factors using *car* package v. 3.1.3 (Fox and Weisberg 2019).

The terminal measurements body fat content and digestive tract organ sizes were modeled separately using the fixed terms line type * diet treatment (including main effects), sex, and the timing of dissection (day 48 or delayed). We also included the dry mass of the bank vole carcass to control for structural size, but this information, which was derived from the fat extraction measurements, was missing for 22 individuals included in the organ data set and was not statistically significant. Therefore, it was kept in the fat model but dropped from the organ model to improve sample size. Replicate line (1|Line) was included as a random intercept in both models. Where the interaction term was non-significant, it was dropped and the model was estimated again with only the additive terms.

To examine longitudinal (within-individual) changes in the microbiota, we used "first distances" for the three different beta diversity metrics as outcome variables in linear mixed models. We computed two separate models for A) the distance from P1 to P2, and B) the distance from P1 to P3. For model A, larger distances are expected for the CR treatment compared to the SD treatment. For model B, we expect similar, minor distances for both treatments if the community has completely recovered. We tested the effect of diet treatment * selection line type (including main effects) on first distances and controlled for sex and clostridium presence. As only one measure per individual was included per model, the random intercept of Line was included as a random effect. When the random effect of Line had a zero variance and caused non-convergence due to singularities, it was removed from the model and a linear model with the same fixed terms was estimated instead (likelihood ratio test of nested models with and without random effect of Line in all cases $X^2$=0, P=1). The overall effect of the interaction term was evaluated by comparing the nested models with anova (likelihood ratio test, models re-fitted with ML instead of REML), and the term was dropped when non-significant.

To identify the taxa that differ in abundance among treatment groups, we used ancombc2 (ANCOMBC v. 2.8.1 (Lin and Peddada 2020, 2024; Lin et al. 2022) with default settings and structural zeros detected based on diet treatment. We initially constructed a model for a three-way interaction of treatment x selection x phase and sex, with random intercept of vole identity nested within line, but this model failed to converge and had to be simplified. Thus, we estimated differential abundances separately for phases 2 and 3 on genus and family level, with each model constructed with the fixed terms treatment, selection, treatment × selection interaction, and sex. Due to consistent singularities resulting from all random effects structures, the random formula was set to NULL for all models. Additionally, we examined the main effect of diet treatment on differentially expressed families at P2 to gain information on the more general effects of dietary change. Bacterial taxa that failed sensitivity analyses for the variable of interest (treatment × selection or treatment) were excluded from the results. Taxa identified as differentially abundant for the interaction term were further examined by modeling the log10-transformed abundance (+1e-6 to account for zeros) of the family/genus within a sample as a function of treatment, selection, treatment × selection, and sex, and the random intercept of line.

### Supplementary results

#### Supplementary tables

Table S1: Predictors of body mass, food consumption, and resting metabolic rate from final models. The significance of the Selection:Treatment:Phase interaction was derived from testing the difference between nested models including and excluding the interaction term. The rest of the results for body mass and food consumption are based on the reduced model after dropping the 3-way interaction term. Statistically significant results in bold.

| | **Resting metabolic rate** | | | | **Body mass** | | | | **Food consumption** | | | |
|---|---|---|---|---|---|---|---|---|---|---|---|---|
| 3-way interaction | LRT | P | | | LRT | P | | | LRT | P | | |
| Selection:Treatment:Phase | 3.603 | 0.165 | | | 1.159 | 0.56 | | | 1.329 | 0.515 | | |
| **Fixed effects** | **Estimate** | **SE** | **t** | **P** | **Estimate** | **SE** | **t** | **P** | **Estimate** | **SE** | **t** | **P** |
| (Intercept) | -3.348 | 0.236 | -14.169 | <0.001 | -0.584 | 0.324 | -1.802 | 0.072 | -0.738 | 0.163 | -4.540 | <0.001 |
| SelectionA | 0.813 | 0.184 | 4.413 | **0.005** | 0.337 | 0.454 | 0.742 | 0.486 | 1.226 | 0.222 | 5.534 | **0.002** |
| TreatmentCR | 0.027 | 0.127 | 0.210 | 0.834 | -0.097 | 0.128 | -0.760 | 0.448 | -0.066 | 0.143 | -0.461 | 0.645 |
| PhaseP2 | -0.172 | 0.099 | -1.747 | 0.081 | 0.101 | 0.050 | 2.033 | **0.043** | -0.065 | 0.093 | -0.701 | 0.484 |
| PhaseP3 | -0.168 | 0.091 | -1.856 | 0.064 | 0.089 | 0.051 | 1.749 | 0.081 | -0.133 | 0.095 | -1.407 | 0.160 |
| Sexmale | 0.031 | 0.075 | 0.416 | 0.678 | 0.894 | 0.085 | 10.496 | **<0.001** | 0.366 | 0.087 | 4.216 | **<0.001** |
| Body mass | 0.112 | 0.010 | 11.552 | **<0.001** | | | | | | | | |
| Activity | 2.118 | 0.144 | 14.662 | **<0.001** | | | | | | | | |
| SelectionA:TreatmentCR | -0.095 | 0.177 | -0.538 | 0.591 | 0.095 | 0.179 | 0.533 | 0.595 | 0.213 | 0.202 | 1.057 | 0.292 |
| SelectionA:PhaseP2 | -0.124 | 0.139 | -0.896 | 0.371 | -0.005 | 0.070 | -0.070 | 0.944 | -0.281 | 0.131 | -2.151 | **0.032** |
| SelectionA:PhaseP3 | -0.238 | 0.128 | -1.864 | 0.063 | 0.086 | 0.071 | 1.211 | 0.227 | -0.232 | 0.132 | -1.751 | 0.081 |
| TreatmentCR:PhaseP2 | 0.058 | 0.145 | 0.401 | 0.689 | -0.376 | 0.071 | -5.266 | **<0.001** | 0.410 | 0.134 | 3.061 | **0.002** |
| TreatmentCR:PhaseP3 | 0.108 | 0.130 | 0.834 | 0.405 | -0.134 | 0.072 | -1.865 | 0.063 | 0.078 | 0.133 | 0.586 | 0.558 |
| SelectionA:TreatmentCR:PhaseP2 | -0.380 | 0.203 | -1.875 | 0.062 | -0.013 | 0.102 | -0.125 | 0.901 | 0.046 | 0.189 | 0.245 | 0.806 |
| SelectionA:TreatmentCR:PhaseP3 | -0.183 | 0.184 | -0.993 | 0.322 | 0.088 | 0.102 | 0.865 | 0.388 | -0.163 | 0.188 | -0.863 | 0.389 |
| **Random effects** | **Std.Dev.** | **N** | **Cor. Str.** | **Var. Str.** | **Std.Dev.** | **N** | **Cor. Str.** | **Var. Str.** | **Std.Dev.** | **N** | **Cor. Str.** | **Var. Str.** |
| ID:Line | 0.085 | 221 | CS Rho | Id Phase | 0.612 | 221 | CS Rho | Fixed | 0.573 | 220 | CS Rho | Id Phase |
| Line | 0.192 | 8 | 0.379 | | 0.617 | 8 | <0.001 | | 0.241 | 8 | -0.199 | |
| Residual | 0.64 | | | | 0.257 | | | | 0.482 | | | |
| N observations | 605 | | | | 630 | | | | 633 | | | |
| $R^2$ marginal | 0.572 | | | | 0.247 | | | | 0.394 | | | |
| $R^2$ conditional | 0.613 | | | | 0.939 | | | | 0.773 | | | |

Table S2: Predictors of body fat content and digestive tract size

| Response: | **Fat content** | | | | **Dry mass of digestive tract organs** | | | |
|---|---|---|---|---|---|---|---|---|
| Treatment:Selection Interaction | $X^2$ 0.582 | Df 1 | P 0.446 | | $X^2$ 2.329 | Df 1 | P 0.127 | |
| **Fixed effects** | **Estimate** | **SE** | **t** | **P** | **Estimate** | **SE** | **t** | **P** |
| (Intercept) | -3.265 | 0.183 | -17.795 | <0.001 | -0.254 | 0.222 | -1.141 | 0.278 |
| TreatmentCR | 0.030 | 0.077 | 0.383 | 0.702 | -0.179 | 0.141 | -1.269 | 0.207 |
| SelectionA | -0.272 | 0.101 | -2.688 | **0.039** | 0.819 | 0.271 | 3.021 | **0.023** |
| Sexmale | -0.092 | 0.078 | -1.175 | 0.242 | -0.002 | 0.137 | -0.011 | 0.991 |
| CarcassDrymass | 0.368 | 0.017 | 21.334 | **<0.001** | | | | |
| DissectionDelayed | 0.070 | 0.100 | 0.698 | 0.486 | -0.256 | 0.171 | -1.498 | 0.136 |
| **Random effects** | **Std.Dev.** | **N** | | | **Std.Dev.** | **N** | | |
| Line | 0.094 | 8 | | | 0.330 | 8 | | |
| Residual | 0.474 | | | | 0.858 | | | |
| N | 161 | | | | 159 | | | |
| $R^2$ marginal | 0.774 | | | | 0.190 | | | |
| $R^2$ conditional | 0.782 | | | | 0.295 | | | |

Table S3: The sixteen most common families (relative abundance >0.01 in unrarefied data) across all treatments and selection lines.

| **Phylum** | **Class** | **Order** | **Family** | **Mean rel. abundance** |
|---|---|---|---|---|
| *Bacteroidota* | *Bacteroidia* | *Bacteroidales* | *Muribaculaceae* | 0.400 |
| *Firmicutes* | *Bacilli* | *Lactobacillales* | *Lactobacillaceae* | 0.170 |
| *Firmicutes* | *Clostridia* | *Lachnospirales* | *Lachnospiraceae* | 0.109 |
| *Firmicutes* | *Bacilli* | *Erysipelotrichales* | *Erysipelotrichaceae* | 0.042 |
| *Firmicutes* | *Clostridia* | *Christensenellales* | *Christensenellaceae* | 0.032 |
| *Bacteroidota* | *Bacteroidia* | *Bacteroidales* | *Rikenellaceae* | 0.026 |
| *Firmicutes* | *Clostridia* | *Oscillospirales* | *Ruminococcaceae* | 0.025 |
| *Firmicutes* | *Clostridia* | *Clostridia_vadinBB60_group* | *Clostridia_vadinBB60_group* | 0.021 |
| *Desulfobacterota* | *Desulfovibrionia* | *Desulfovibrionales* | *Desulfovibrionaceae* | 0.020 |
| *Campilobacterota* | *Campylobacteria* | *Campylobacterales* | *Helicobacteraceae* | 0.019 |
| *Bacteroidota* | *Bacteroidia* | *Bacteroidales* | *Rs-E47_termite_group* | 0.018 |
| *Firmicutes* | *Clostridia* | *Oscillospirales* | *Oscillospiraceae* | 0.016 |
| *Spirochaetota* | *Spirochaetia* | *Spirochaetales* | *Spirochaetaceae* | 0.014 |
| *Actinobacteriota* | *Actinobacteria* | *Bifidobacteriales* | *Bifidobacteriaceae* | 0.010 |
| *Fusobacteriota* | *Fusobacteriia* | *Fusobacteriales* | *Fusobacteriaceae* | 0.010 |

Table S4: Longitudinal changes (between P1, P2, and P3) in gut microbiota alpha diversity (Observed ASVs and Shannon index) in selection lines of the bank vole in response to a change in diet. The overall effect of the three-way interaction Selection:Treatment:Phase was estimated by comparing nested models including and excluding the term.

| | **Observed ASVs** | | | | **Shannon index** | | | |
|---|---|---|---|---|---|---|---|---|
| 3-way interaction | $X^2$ | df | P | | $X^2$ | df | P | |
| Selection:Treatment:Phase | 8.720 | 2 | **0.013** | | 8.581 | 2 | **0.014** | |
| **Fixed effects** | **Estimate** | **SE** | **z** | **P** | **Estimate** | **SE** | **t** | **P** |
| (Intercept) | 5.889 | 0.086 | 68.750 | <0.001 | 3.739 | 0.106 | 35.317 | <0.001 |
| SelectionA | -0.215 | 0.113 | -1.910 | 0.056 | -0.193 | 0.143 | -1.349 | 0.194 |
| TreatmentCR | -0.187 | 0.100 | -1.880 | 0.060 | -0.232 | 0.114 | -2.040 | **0.042** |
| PhaseP2 | 0.007 | 0.048 | 0.150 | 0.880 | 0.069 | 0.084 | 0.823 | 0.411 |
| PhaseP3 | -0.023 | 0.047 | -0.480 | 0.634 | 0.094 | 0.083 | 1.133 | 0.258 |
| Sexmale | -0.065 | 0.065 | -0.990 | 0.322 | -0.075 | 0.064 | -1.177 | 0.241 |
| SelectionA:TreatmentCR | 0.255 | 0.142 | 1.800 | 0.072 | 0.287 | 0.161 | 1.783 | 0.076 |
| SelectionA:PhaseP2 | 0.084 | 0.068 | 1.240 | 0.215 | 0.223 | 0.119 | 1.876 | 0.062 |
| SelectionA:PhaseP3 | 0.104 | 0.067 | 1.540 | 0.125 | 0.077 | 0.118 | 0.647 | 0.518 |
| TreatmentCR:PhaseP2 | 0.151 | 0.068 | 2.220 | **0.026** | 0.794 | 0.119 | 6.665 | **<0.001** |
| TreatmentCR:PhaseP3 | 0.039 | 0.069 | 0.570 | 0.568 | 0.159 | 0.120 | 1.323 | 0.187 |
| SelectionA:TreatmentCR:PhaseP2 | -0.237 | 0.096 | -2.470 | **0.013** | -0.442 | 0.168 | -2.629 | **0.009** |
| SelectionA:TreatmentCR:PhaseP3 | -0.258 | 0.097 | -2.660 | **0.008** | -0.404 | 0.169 | -2.390 | **0.017** |
| **Random effects** | **Std.Dev.** | **N** | **Dispersion** | | **Std.Dev.** | **N** | | |
| Intercept ID:Line | 0.402 | 168 | 24.6 | | 0.348 | 168 | | |
| Intercept Line | 0.072 | 8 | | | 0.123 | 8 | | |
| N observations | 474 | | | | 474 | | | |
| $R^2$ marginal | 0.04 * | | | | 0.02 | | | |
| $R^2$ conditional | 0.80 * | | | | 0.60 | | | |

* pseudo-$R^2$ based on delta, lognormal and trigamma approximation

Table S5: Models of first distances (log-transformed) by three different beta diversity metrics. First panel P1-P2 reflects magnitude of within-individual change in bacterial community from baseline P1 to the end of diet treatment P2. The second panel P1-P3 reflects the completeness of recovery, from baseline P1 to the end of the recovery period P3. SS= sum of squares.

| Bray-Curtis | **P1-P2 first distance** | | | | **P1-P3 first distance** | | | |
|---|---|---|---|---|---|---|---|---|
| Selection:Treatment interaction | $X^2$ | Df | P | | $X^2$ | Df | P | |
| | 0.076 | 1 | 0.782 | | <0.001 | 1 | 0.985 | |
| **Fixed effects** | **Estimate** | **SE** | **t** | **P** | **Estimate** | **SE** | **t** | **P** |
| (Intercept) | 0.353 | 0.014 | 25.687 | <0.001 | 0.409 | 0.024 | 17.300 | **<0.001** |
| TreatmentCR | 0.153 | 0.012 | 13.187 | **<0.001** | 0.064 | 0.015 | 4.207 | **<0.001** |
| SelectionA | 0.003 | 0.015 | 0.165 | 0.875 | -0.042 | 0.030 | -1.391 | 0.213 |
| Sexmale | 0.014 | 0.012 | 1.245 | 0.215 | -0.028 | 0.015 | -1.816 | 0.072 |
| **Random effects** | **Std.Dev.** | **N** | | | **Std.Dev.** | **N** | | |
| Line | 0.014 | 8 | | | 0.036 | 8 | | |
| Residual | 0.071 | | | | 0.092 | | | |
| N | 152 | | | | 147 | | | |
| $R^2$ marginal | 0.527 | | | | 0.142 | | | |
| $R^2$ conditional | 0.545 | | | | 0.258 | | | |
| **Unweighted Unifrac** | **P1-P2 first distance** | | | | **P1-P3 first distance** | | | |
| Selection:Treatment interaction | SS | F | P | | SS | F | P | |
| | -0.001 | 0.139 | 0.710 | | -0.002 | 0.312 | 0.578 | |
| **Fixed effects** | **Estimate** | **SE** | **t** | **P** | **Estimate** | **SE** | **t** | **P** |
| (Intercept) | 0.289 | 0.010 | 27.801 | <0.001 | 0.301 | 0.013 | 24.067 | <0.001 |
| TreatmentCR | 0.027 | 0.010 | 2.642 | **0.009** | 0.020 | 0.013 | 1.571 | 0.118 |
| SelectionA | 0.001 | 0.010 | 0.083 | 0.934 | -0.004 | 0.013 | -0.294 | 0.770 |
| Sexmale | -0.011 | 0.010 | -1.021 | 0.309 | 0.000 | 0.013 | -0.016 | 0.987 |
| N | 148 | | | | 144 | | | |
| $R^2$ | 0.052 | | | | 0.017 | | | |
| $R^2$ adj. | 0.033 | | | | -0.003 | | | |
| **Weighted Unifrac** | **P1-P2 first distance** | | | | **P1-P3 first distance** | | | |
| Selection:Treatment interaction | SS | F | P | | $X^2$ | Df | P | |
| | <-0.001 | 0.318 | 0.573 | | 0.137 | 1 | 0.711 | |
| **Fixed effects** | **Estimate** | **SE** | **t** | **P** | **Estimate** | **SE** | **t** | **P** |
| (Intercept) | 0.132 | 0.008 | 16.590 | <0.001 | 0.158 | 0.013 | 12.244 | 0.000 |
| TreatmentCR | 0.084 | 0.008 | 10.702 | **0.007** | 0.024 | 0.009 | 2.703 | **0.008** |
| SelectionA | -0.009 | 0.008 | -1.152 | 0.251 | -0.027 | 0.016 | -1.675 | 0.144 |
| Sexmale | 0.003 | 0.008 | 0.427 | 0.670 | -0.015 | 0.009 | -1.671 | 0.097 |
| **Random effects** | | | | | **Std.Dev.** | **N** | | |
| Line | | | | | 0.019 | 8 | | |
| Residual | | | | | 0.054 | | | |
| N | 148 | | | | 147 | | | |
| $R^2$ unadjusted/marginal | 0.438 | | | | 0.103 | | | |
| $R^2$ adjusted/conditional | 0.427 | | | | 0.199 | | | |

Table S6: Model predicted values for *Helicobacteraceae* relative abundance (log10(abundance + 1e-6)) at the end of dietary restriction (phase P2).

| Fixed effects | Estimate | SE | df | t | P |
|---|---|---|---|---|---|
| (Intercept) | -2.560 | 0.176 | 27.970 | -14.521 | <0.001 |
| TreatmentCR | 0.821 | 0.210 | 147.955 | 3.907 | <0.001 |
| SelectionA | 0.587 | 0.225 | 19.097 | 2.613 | 0.017 |
| Sexmale | -0.042 | 0.149 | 148.018 | -0.279 | 0.781 |
| TreatmentCR:SelectionA | -0.863 | 0.298 | 147.938 | -2.896 | 0.004 |
| Random effects | Variance | Std.Dev. | N | | |
| Line | 0.011 | 0.103 | 8 | | |
| Residual | 0.882 | 0.939 | | | |
| N | 159 | | | | |
| $R^2$ marginal | 0.093 | | | | |
| $R^2$ conditional | 0.104 | | | | |

Table S7: Differentially abundant families for diet treatments at phases P2 and P3. Enrichment in CR treatment is indicated by positive log fold change (LFC), negative LFC indicates enrichment in SD treatment.

| Phase P2 | | | | | | | | |
|---|---|---|---|---|---|---|---|---|
| **Phylum** | **Class** | **Order** | **Family** | **LFC** | **SE** | **W** | **$P_{fdr}$** | **Mean rel. abundance** |
| *Proteobacteria* | *Gammaproteobacteria* | *Burkholderiales* | *Oxalobacteraceae* | 0.892 | 0.130 | 6.834 | <0.001 | 0.0003 |
| *Actinobacteriota* | *Coriobacteriia* | *Coriobacteriales* | *Eggerthellaceae* | 0.724 | 0.143 | 5.055 | <0.001 | 0.0029 |
| *Firmicutes* | *Clostridia* | *Lachnospirales* | *Lachnospiraceae* | 0.549 | 0.096 | 5.730 | <0.001 | 0.1093 |
| *Firmicutes* | *Clostridia* | *Oscillospirales* | *[Clostridium]_methylpentosum_group* | -0.457 | 0.098 | -4.661 | 0.010 | <0.0001 |
| *Verrucomicrobiota* | *Verrucomicrobiae* | *Opitutales* | *Puniceicoccaceae* | -0.757 | 0.146 | -5.195 | <0.001 | 0.0004 |
| *Proteobacteria* | *Alphaproteobacteria* | *Paracaedibacterales* | *Paracaedibacteraceae* | -0.810 | 0.196 | -4.127 | 0.002 | 0.0021 |
| *Firmicutes* | *Clostridia* | *Clostridia_UCG-014* | *Clostridia_UCG-014* | -0.852 | 0.178 | -4.786 | <0.001 | 0.0018 |
| *Spirochaetota* | *Spirochaetia* | *Spirochaetales* | *Spirochaetaceae* | -0.978 | 0.184 | -5.313 | <0.001 | 0.0144 |
| *Cyanobacteria* | *Vampirivibrionia* | *Gastranaerophilales* | *Gastranaerophilales* | -1.297 | 0.212 | -6.108 | <0.001 | 0.0069 |
| *Firmicutes* | *Bacilli* | *Lactobacillales* | *Lactobacillaceae* | -1.903 | 0.226 | -8.434 | <0.001 | 0.1704 |
| *Firmicutes* | *Bacilli* | *Erysipelotrichales* | *Erysipelotrichaceae* | -2.090 | 0.410 | -5.103 | <0.001 | 0.0428 |
| *Actinobacteriota* | *Actinobacteria* | *Bifidobacteriales* | *Bifidobacteriaceae* | -3.617 | 0.224 | -16.136 | <0.001 | 0.0105 |
| *Firmicutes* | *Clostridia* | *Christensenellales* | *Christensenellaceae* | -4.501 | 0.295 | -15.262 | <0.001 | 0.0317 |
| *Phase P3* | | | | | | | | |
| *Firmicutes* | *Clostridia* | *Christensenellales* | *Christensenellaceae* | -2.541 | 0.337 | -7.538 | <0.001 | 0.0317 |
| *Firmicutes* | *Clostridia* | *Clostridia_UCG-014* | *Clostridia_UCG-014* | -0.691 | 0.177 | -3.910 | 0.007 | 0.0018 |

#### Supplementary figures

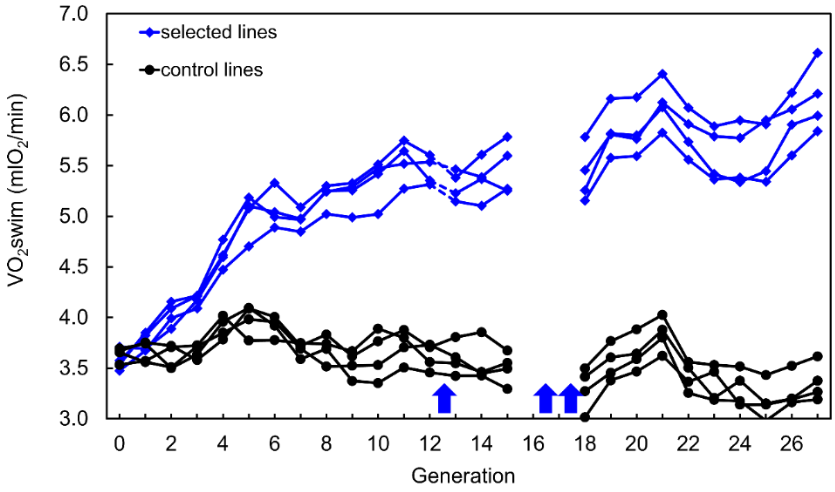

Figure S1. Differences in oxygen consumption ($V0_2$) between A and C lines during swimming trials.

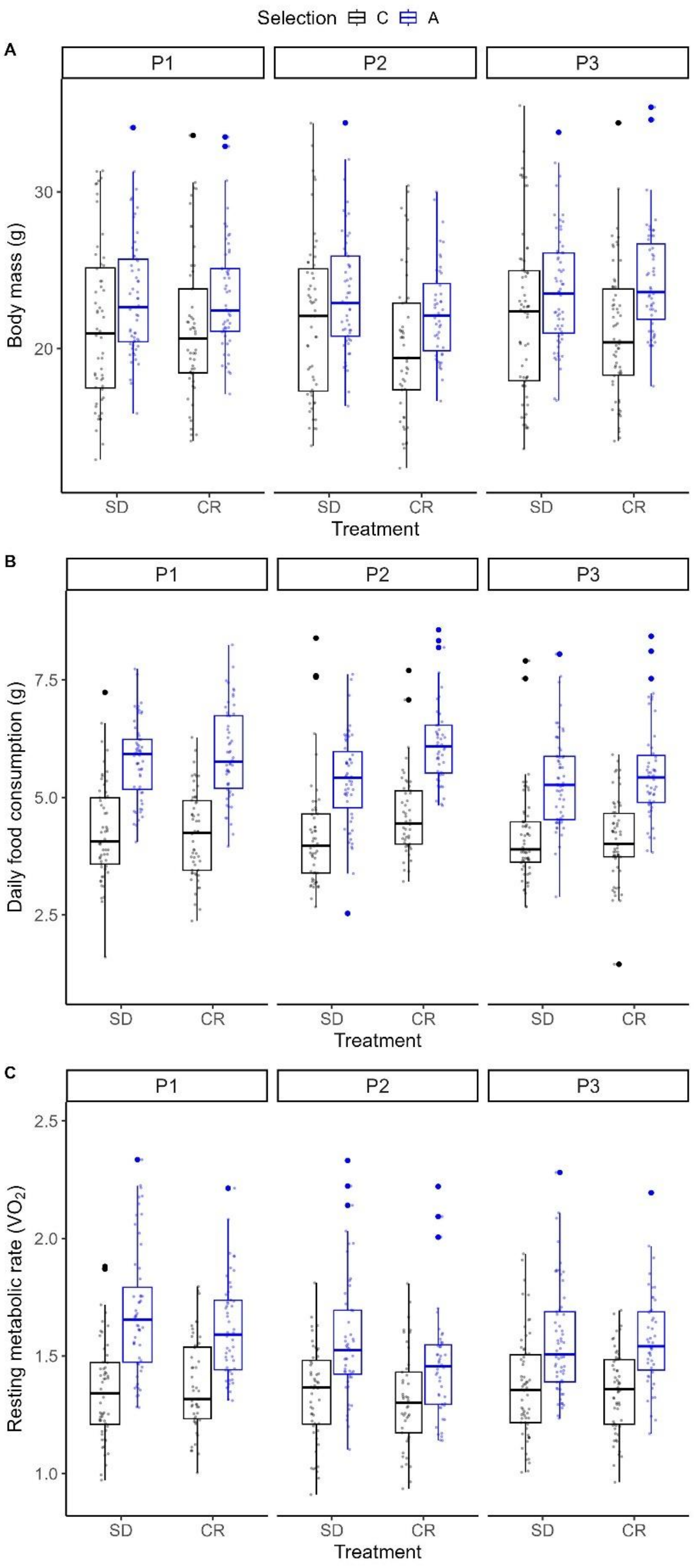

Figure S2: Raw data for A) body mass, B) daily food consumption, and 3) RMR (average of minimum 3 measurements) across the three phases of the experiment.

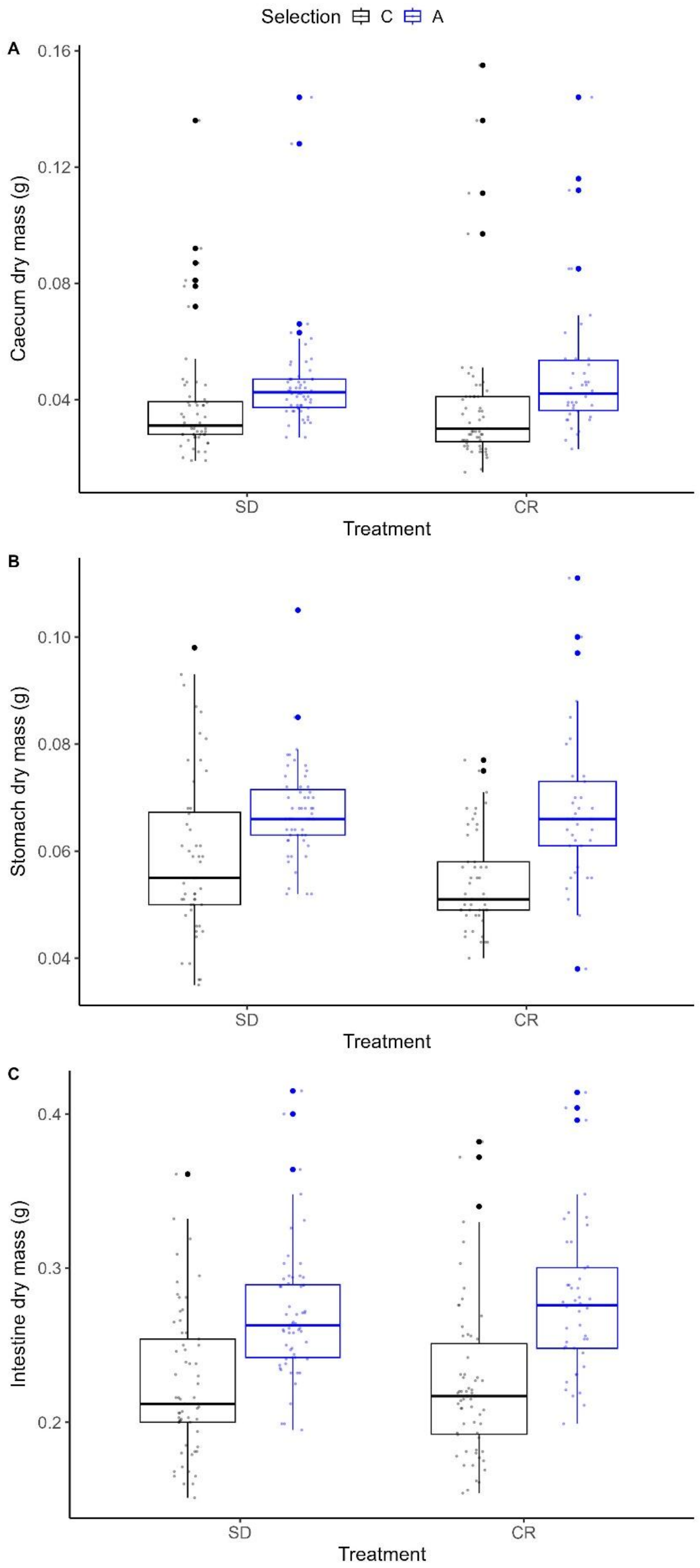

Figure S3. Digestive organ (A: caecum, B: stomach, C: small and large intestine) dry mass in the two food treatments and selection regimes (raw data excluding outliers ±3 SD).

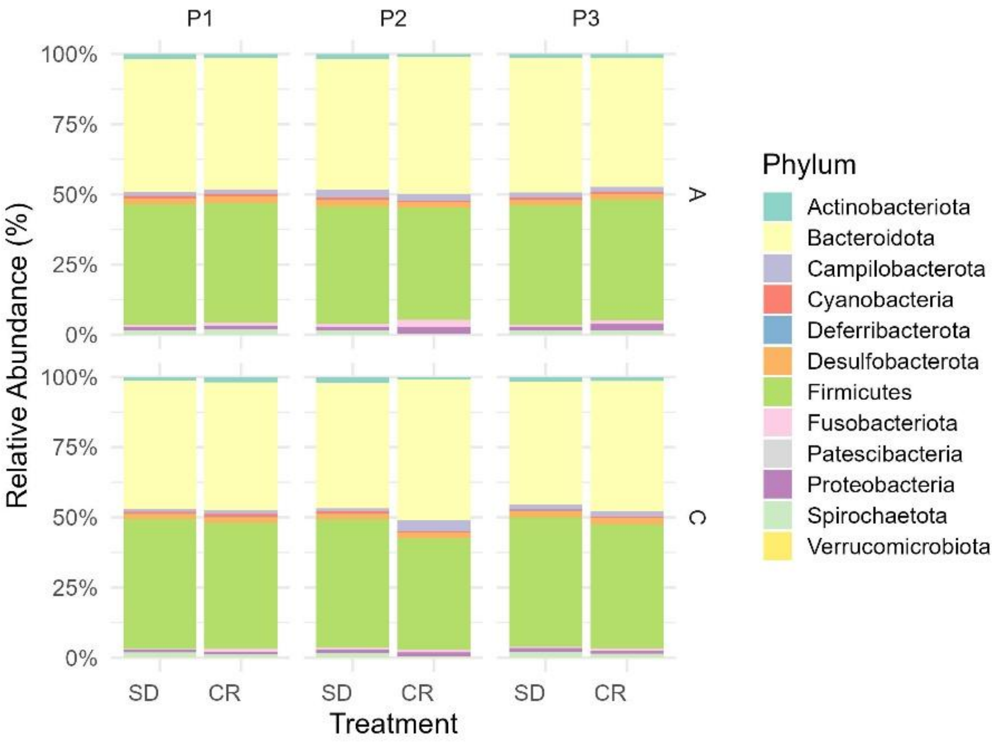

Figure S4: Relative abundances of phyla in the three experiment phases in SD and CR treatments. Top row shows data for A-lines, bottom row C-lines.

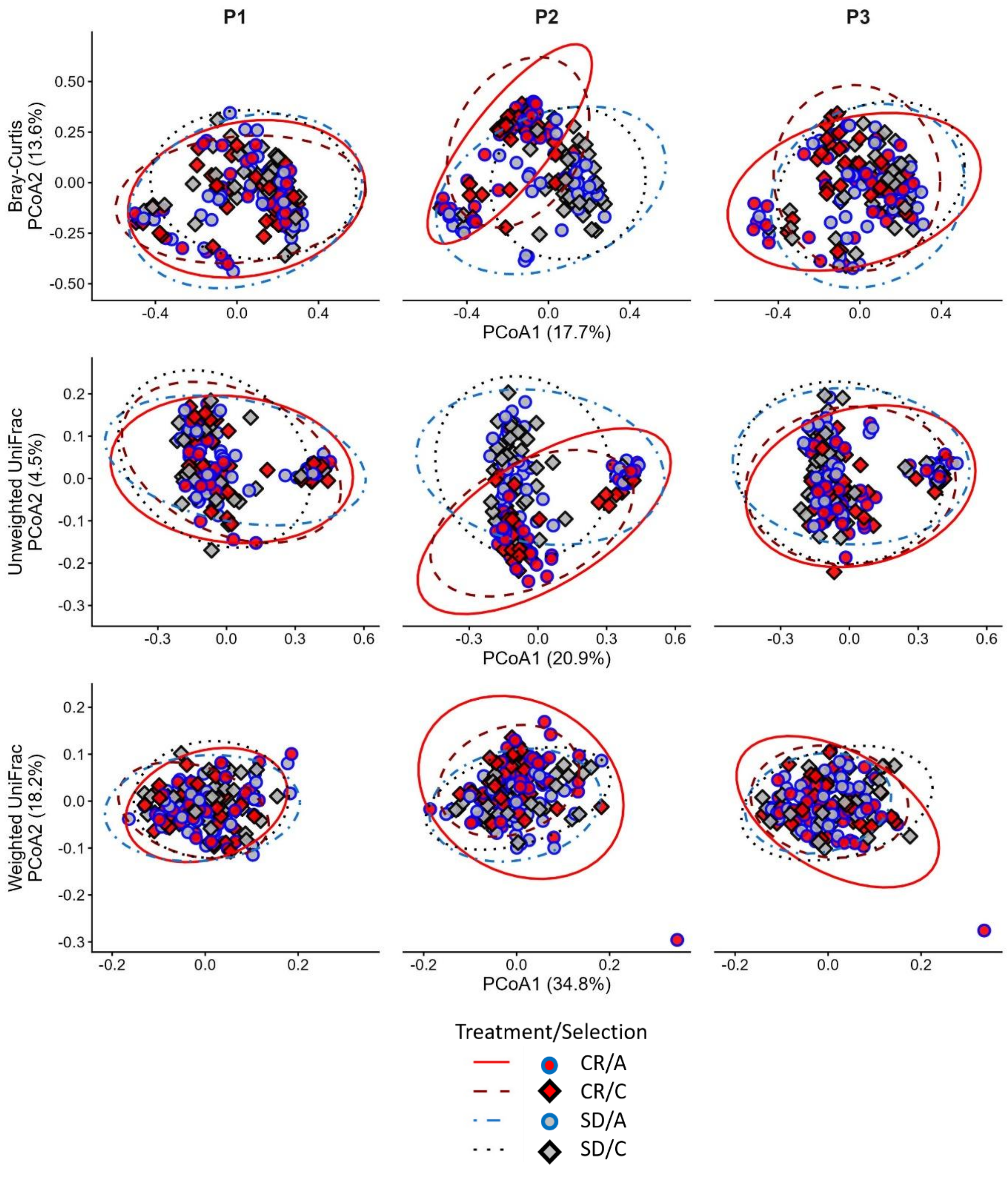

Fig. S5. Principal coordinate analysis (PCoA) ordination based on A) Bray-Curtis, B) Unweighted UniFrac, and C) Weighted UniFrac dissimilarity matrices showing community composition of bank vole gut microbiota among diet treatments (SD, CR) and selection line types (C, A) at the three experimental phases (P1 baseline, P2 end of diet treatment, P3 end of recovery).

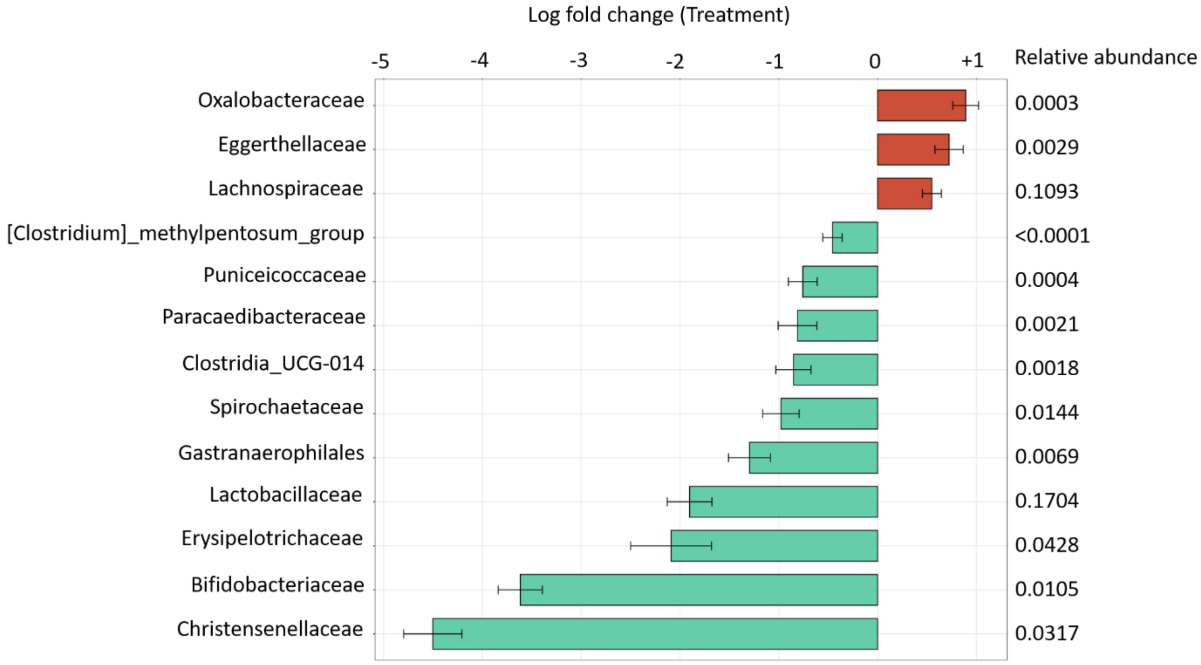

Figure S6: Differentially abundant bacterial families between dietary treatments at P2. Positive log fold change (orange) shows overrepresented families in CR treatment over SD, while negative log fold change (blue) indicates overrepresentation of the family in SD relative to CR treatment. Whiskers indicate standard errors. Only robust findings (passed sensitivity analysis for treatment term) from bias-corrected analyses at $P_{fdr}<0.05$ are shown.
